# Full-length transcriptomics and proteomics reveal how genome minimization reshapes gene expression in synthetic bacteria

**DOI:** 10.64898/2026.09.08.750226

**Authors:** Enguang Fu, Troy A. Brier, Jay E. Cournoyer, Sage A. Glass, Yang-le Gao, Yanbao Yu, Zane R. Thornburg, Karrie Goglin, Gavin John, Tiana Mamaghani, Sujata Shivakumar, Abner T. Apsley, Christopher J. Fields, Angad P. Mehta, John I. Glass, Zaida Luthey-Schulten

## Abstract

JCVI-syn1.0 (Syn1.0) and JCVI-syn3A (Syn3A), genetically synthetic and genome-reduced versions of the naturally occurring bacterium *Mycoplasma mycoides*, are landmark platforms for defining the gene set required for life, yet how their genomes are expressed at the RNA level remains uncharacterized. We combined full-length PacBio and native long-read RNA sequencing with short-read quantification and complementary proteomics to map transcription, RNA processing, and protein abundance in both cells. In Syn1.0, full-length sequencing resolved 459 operons encompassing 911 genes, revealing pervasive RNA processing with a strong 3*^′^* bias. Analysis of the division and cell-wall cluster showed how transcriptional context explained the restoration of genes required for normal cell division in Syn3A. Most antisense and intergenic transcription in Syn1.0 reflected low-level transcriptional noise arising from inherited mis-annotation, read-through, and synthetic sequences. Much of that transcription was lost in Syn3A after genome minimization. Reducing the genome unexpectedly altered the expression of several retained genes by deleting promoters, most prominently reducing expression of the nucleoid protein HupA and central-carbon enzymes. Meanwhile, one third of the coding mRNA pool was allocated to a single 21-gene ribosomal-protein operon, while several other ribosomal proteins had reduced transcript abundance, which possibly led to imbalanced ribosome assembly. Expression of RNA polymerase and central-carbon metabolism declined at both mRNA and protein levels whereas RNase Y degradosome abundance increased. These shifts in the synthetic evolution of Syn3A suggest plausible mechanisms for its reduced chromosome contacts and slower growth. Together, these results show that genome minimization alters not only gene content but also the transcriptional context and resource allocation of retained genes. The analysis and visualization that are shared via Jupyter Notebook provide the RNA-level foundation for whole-cell modeling of the minimal cell.

## Introduction

Genetically minimized bacteria provide a unique opportunity to elucidate the inherent phenomena governing bacterial life [1, 2]. JCVI-syn1.0 (Syn1.0, GenBank CP002027.1) and its minimized derivative JCVI-syn3A (Syn3A, GenBank CP016816.2) are genetically modified and reduced versions of the naturally occurring bacterium *Mycoplasma mycoides* (*M. mycoides*)[3, 4] and they are landmark platforms for defining the minimal requirements of cellular life. In synthesizing the Syn1.0 genome (1,078,809 bp, 911 genes), 14 wild-type *M. mycoides* genes were either deleted or disrupted and five non-native genes required for the cloning procedure in yeast as well as four watermarks and bacterial genes *tetM* and *lacZ* were incorporated [3]. Genome reduction resulted in Syn3.0, which to date has the least number of genes in any bacteria [4]. However, its inconsistent morphology motivated the re-introduction of 19 genes resulting in more robust Syn3A that has normal cell division [2, 5, 6]. Reducing the genome from Syn1.0 to Syn3A (543,379 bp, 493 genes), while preserving the ancestral order of the retained genes, produced the minimal genome of an autonomously replicating organism. An important derivative, Syn3B was created by the introduction of a landing pad into Syn3A genome to facilitate the easy insertion, knockout and testing of new genes [7]. Syn3A and Syn3B have drawn interest from laboratories worldwide seeking the essence of life, through studies of gene essentiality and metabolism [2], three-dimensional organization through cryo-electron tomography [8], cell division [5], lipidomics [9] and in-lab evolution [10, 11]. Together, these efforts enabled the first whole-cell model (WCM) resolved in three-dimensional space across an entire cell cycle [6, 12].

The transcriptional regulatory architecture of Syn1.0 and Syn3A is simple, making the underlying mechanisms easier to resolve. *Mycoplasmas* as obligate parasites living in stable host niches are themselves a product of natural genome reduction. They have descended from low-G+C Gram-positive ancestors through massive gene loss [13]. Syn1.0 (same as *M. mycoides*) retains only a single sigma factor and a small set of transcriptional regulators (18 genes), alongside a handful of housekeeping non-coding RNAs (Table S1). Syn3A carries the reduction further, but by design rather than by evolution: its genome was engineered to keep only the genes needed for growth in a constant laboratory medium [4, 2], and it discards 6 of the prior 18 regulators while keeping the same lone sigma factor. With few dedicated regulators and minimal regulatory RNAs, transcription in Syn1.0 and Syn3A is largely constitutive.

**Table S1:**
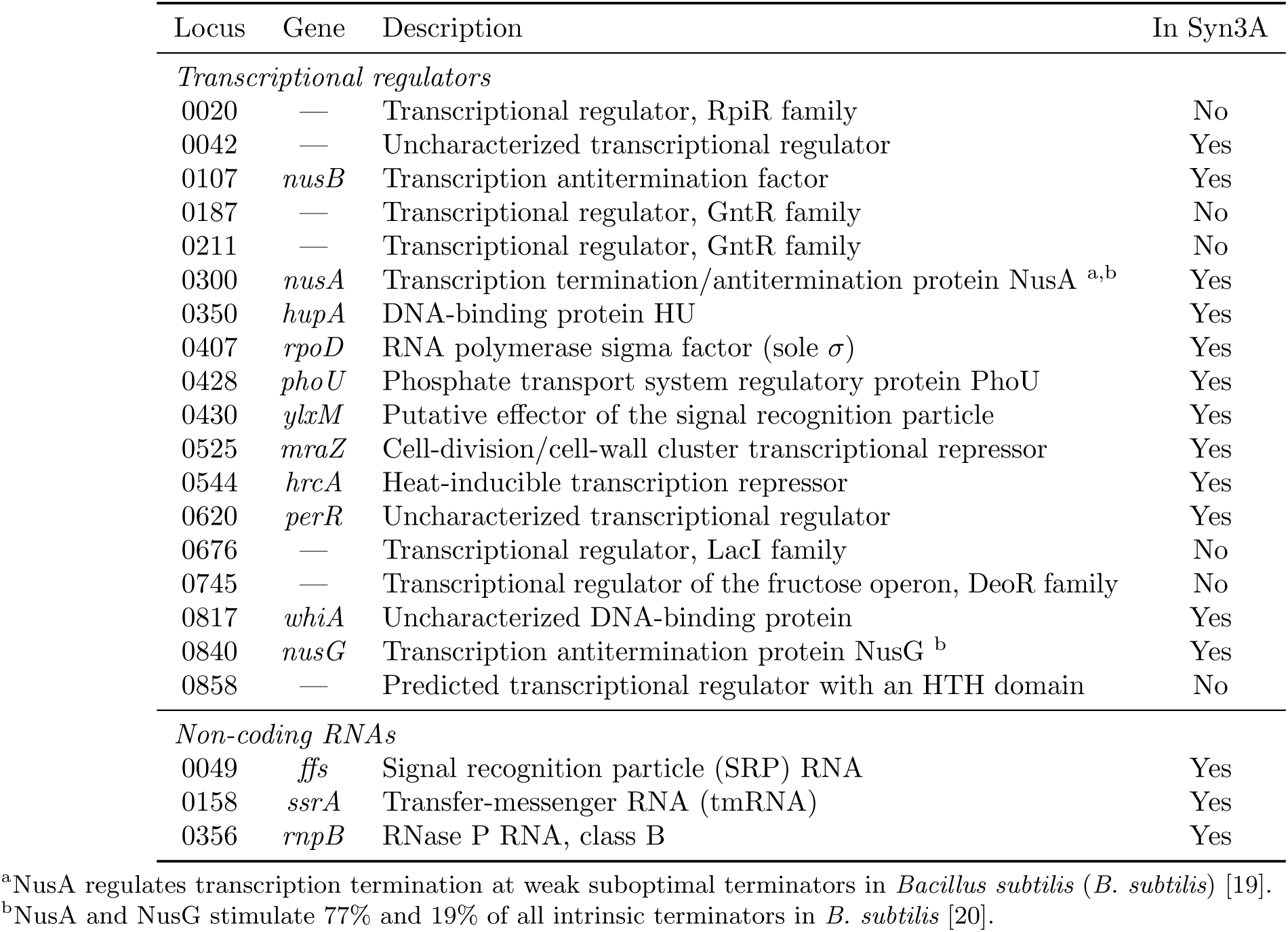
Transcriptional regulators and non-coding RNAs of Syn1.0 and Syn3A. Protein-coding transcriptional regulators (from GenBank functional classification and [2]) and structural or housekeeping non-coding RNAs. Locus numbers are shared between the two organisms; the “In Syn3A” column marks retention after minimization.

| Locus | Gene | Description | In Syn3A |
| --- | --- | --- | --- |
| *Transcriptional regulators* |  |  |  |
| 0020 | — | Transcriptional regulator, RpiR family | No |
| 0042 | — | Uncharacterized transcriptional regulator | Yes |
| 0107 | *nusB* | Transcription antitermination factor | Yes |
| 0187 | — | Transcriptional regulator, GntR family | No |
| 0211 | — | Transcriptional regulator, GntR family | No |
| 0300 | *nusA* | Transcription termination/antitermination protein NusA <sup>a,b</sup> | Yes |
| 0350 | *hupA* | DNA-binding protein HU | Yes |
| 0407 | *rpoD* | RNA polymerase sigma factor (sole $\sigma$) | Yes |
| 0428 | *phoU* | Phosphate transport system regulatory protein PhoU | Yes |
| 0430 | *ylxM* | Putative effector of the signal recognition particle | Yes |
| 0525 | *mraZ* | Cell-division/cell-wall cluster transcriptional repressor | Yes |
| 0544 | *hrcA* | Heat-inducible transcription repressor | Yes |
| 0620 | *perR* | Uncharacterized transcriptional regulator | Yes |
| 0676 | — | Transcriptional regulator, LacI family | No |
| 0745 | — | Transcriptional regulator of the fructose operon, DeoR family | No |
| 0817 | *whiA* | Uncharacterized DNA-binding protein | Yes |
| 0840 | *nusG* | Transcription antitermination protein NusG <sup>b</sup> | Yes |
| 0858 | — | Predicted transcriptional regulator with an HTH domain | No |
| *Non-coding RNAs* |  |  |  |
| 0049 | *ffs* | Signal recognition particle (SRP) RNA | Yes |
| 0158 | *ssrA* | Transfer-messenger RNA (tmRNA) | Yes |
| 0356 | *rnpB* | RNase P RNA, class B | Yes |

<sup>a</sup>NusA regulates transcription termination at weak suboptimal terminators in *Bacillus subtilis* (*B. subtilis*) [19].
<sup>b</sup>NusA and NusG stimulate 77% and 19% of all intrinsic terminators in *B. subtilis* [20].

Many bacterial genes are co-transcribed in operons [14], with multiple transcription start and termination sites (TSS and TTS) acting across a single gene cluster, and downstream endo- and exoribonucleolytic processing further generates a spectrum of overlapping and truncated RNA isoforms (Fig. 1a). Resolving operons requires reading transcripts at full length: short-read Illumina sequencing quantifies transcript abundance robustly but fragments RNA and cannot show which genes share a transcript [15], whereas long-read sequencing PacBio and Oxford Nanopore Technologies (ONT) recover full-length molecules and so reveals operon co-transcription, isoform processing, and antisense or intergenic transcription [16, 17]. A recent custom full-length RNA sequencing protocol, SEnd-seq, revealed unexpected bidirectional transcription terminations [18]. See Methods for detailed comparison of long-read and short-read RNA sequencing platforms.

**Figure 1:**
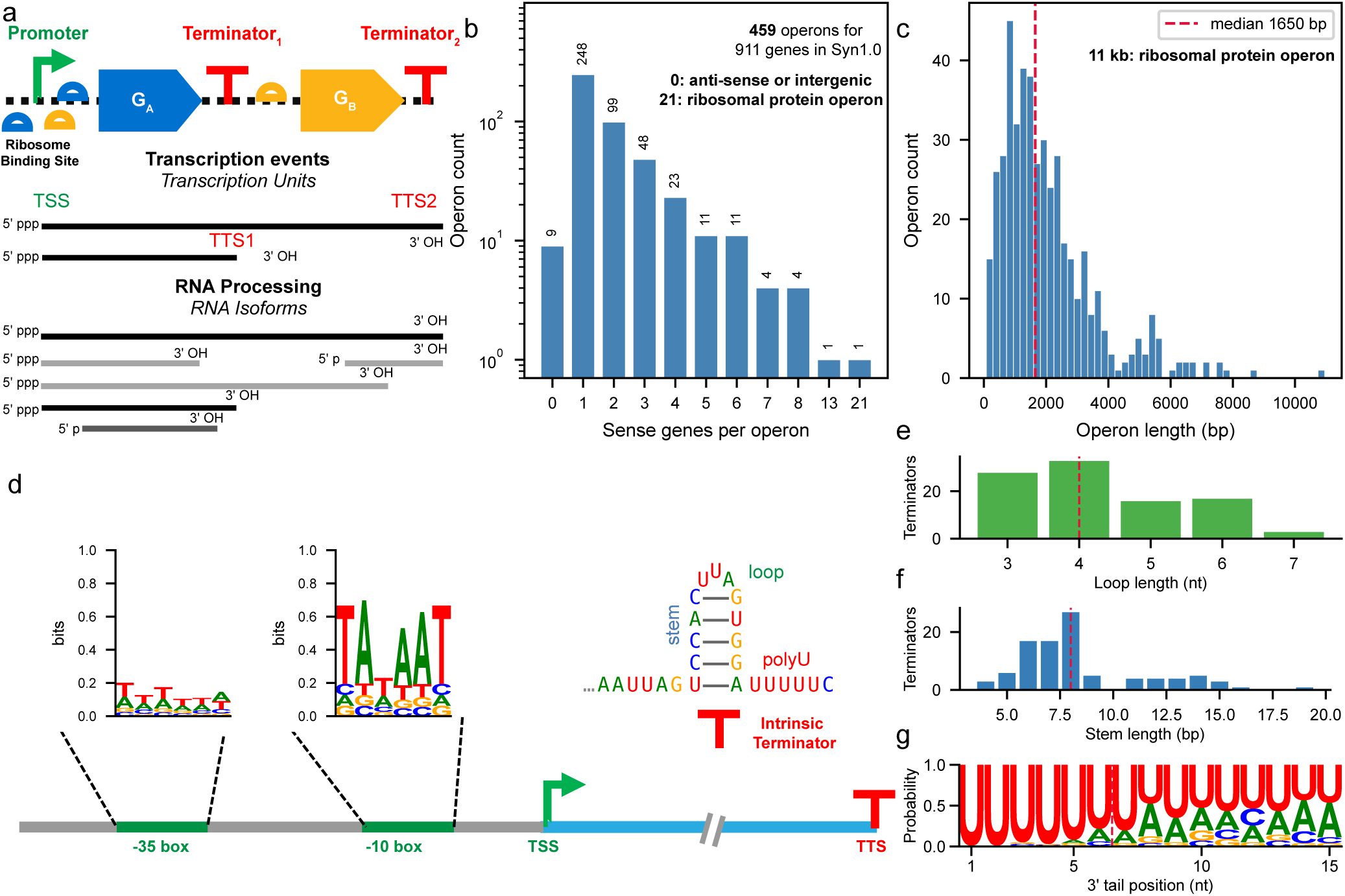
Operon architecture of Syn1.0 from PacBio long-read RNA sequencing. (a) Schematic of two-gene operon co-transcription and subsequent RNA processing into isoforms. (b) Sense genes per operon. (c) Operon length distribution. (d) Promoter and terminator signatures: *−*35 and *−*10 sequence logos at canonical TSSs, and the intrinsic terminator of the *dnaA*/0001–*dnaN* /0002 operon. (e) Terminator loop-length distribution. (f) Terminator stem-length distribution. (g) Terminator 3*^′^* poly-U tail sequence logo. Longest observed polyU length is 15 nt.

Despite the importance of operonal structures in bacterial gene expression, both Syn1.0 and Syn3A have so far been defined chiefly at the level of genome design and protein essentiality [2], leaving how their genes are organized at the RNA level largely uncharacterized. The two also differ in phenotype: Syn3A divides more slowly (2 hours instead of 1 hour for Syn1.0 [2, 5]), packs a denser cytosol [8], and lacks persistent chromosome supercoiling, the last linked to its collapsed level of the nucleoid associated protein HupA [8]. Here, we address three questions that have emerged in the gene expression of Syn1.0 and Syn3A. First, how is gene expression organized across a synthetic genome, from operon boundaries and transcription signals to downstream RNA processing? Second, beyond the deletions themselves, what does this minimization, as a form of synthetic evolution, do to the transcription and translation of the retained genes? And finally, how does that minimization contribute to the phenotypic differences between the two cells? We quantitatively addressed these questions by capturing full-length transcripts with PacBio and ONT in Syn1.0 and ONT in Syn3A, and used Illumina short reads as the standard for transcript quantification. We then combined these with proteomics to build a genome-wide, operon-resolved map of transcription, RNA processing, and translation in both cells. This atlas supplies the RNA-level foundation of gene co-expression and quantification that whole-cell modeling of the minimal cell has so far lacked.

## Results

### Comparison of the multi-platform RNA sequencing of the Syn1 and Syn3A transcriptomes

The Syn1.0 and Syn3A transcriptomes were sequenced on complementary platforms whose depth, read length, and cross-platform agreement were summarized in Table S2 and Fig. S1. In Fig. S1a for Syn1.0, as PacBio Iso-Seq showed the greatest sequencing depth (4,184*×* genome-mean) with the longest reads (median 1.56 kb), several-fold longer than the Syn1.0 ONT direct-RNA reads (median 338–458 nt), the PacBio isoforms were selected to identify operons. It is worth mentioning that transcripts shorter than 200 nt were removed during size selection in the PacBio Iso-seq library preparation, which for example, biased sequencing of *rpmF* /0526 in Fig. 2.

**Figure 2:**
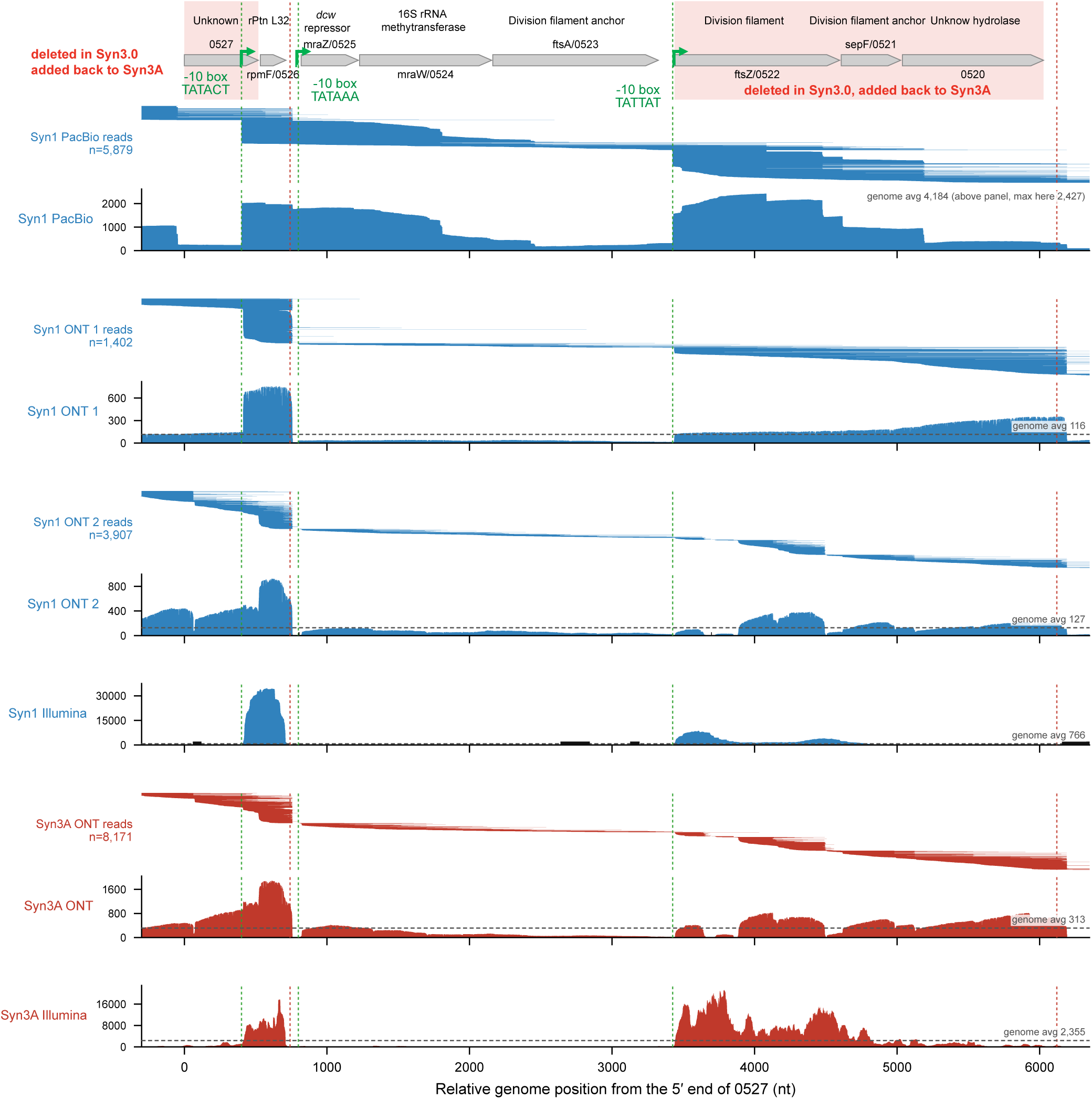
Fine transcriptional structure of the *dcw* cluster (*mraZ* /0525–MMSYN1_0520) and its upstream genes *rpmF* /0526 and MMSYN1_0527, Syn1.0 versus Syn3A. Sense (operon-strand) sequencing depth across the cluster with absolute axis for all six libraries: Syn1.0 PacBio, the two Syn1.0 ONT runs, Syn1.0 Illumina, and the Syn3A ONT and Illumina [10] libraries. Above each long-read library depth, all reads in this region were drawn as horizontal lines ordered by 5*^′^* end. The x-axis is the relative position from the 5*^′^* end of MMSYN1_0527 (both organisms anchored at their own MMSYN1_0527). Green dashed lines mark transcription start sites (a strong promoter inside MMSYN1_0527, a weak internal promoter at *mraZ* /0525, and the *ftsZ* /0522 promoter); red dashed lines mark TransTermHP intrinsic terminators (after *rpmF* /0526 and after MMSYN1_0520). Red bands mark MMSYN1_0527 and the *ftsZ* /0522–MMSYN1_0520 block, deleted in JCVI-syn3.0 but restored in Syn3A.

**Figure S1:**
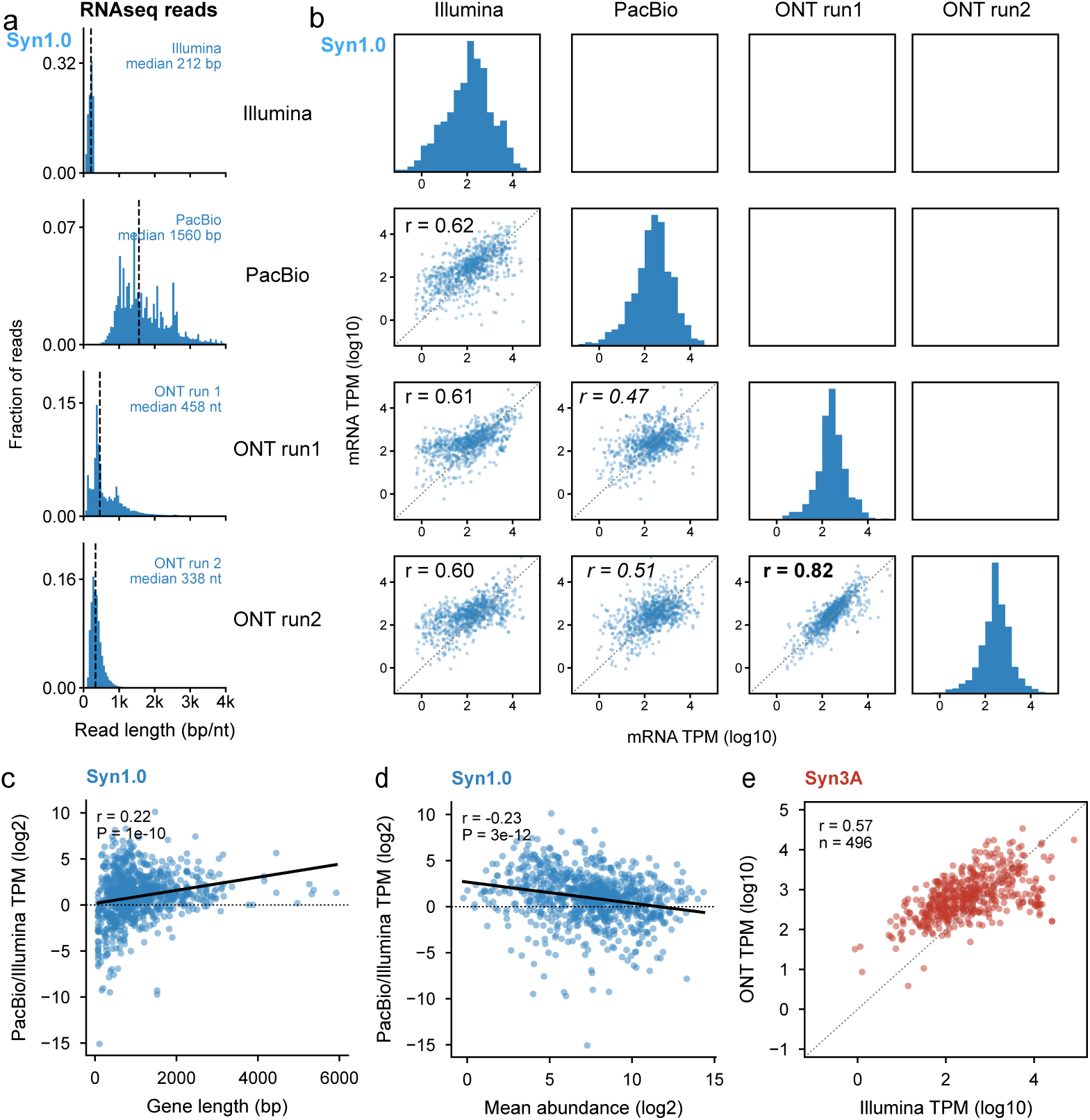
Cross-platform and cross-run comparison of RNA-seq quantification. **(a)** Syn1.0 mapped- read length distribution for each platform (Illumina, PacBio, ONT run 1, ONT run 2; top to bottom) on a shared axis, with the median marked; short-read Illumina, intermediate ONT, and long-read PacBio occupy distinct length regimes. **(b)** Syn1.0 per-gene mRNA TPM across Illumina, PacBio, and the two ONT runs, in that order along both axes: lower triangle, pairwise log_10_–log_10_ scatter with the identity line; diagonal, per-platform log_10_ TPM distribution. **(c)** Syn1.0 gene-length dependence of the PacBio-to-Illumina TPM ratio, with a linear fit. **(d)** The same PacBio-to-Illumina ratio against mean of log_2_ abundance of PacBio and Illumina. **(e)** Syn3A ONT versus Illumina [10] TPM (log_10_–log_10_).

**Table S2:**
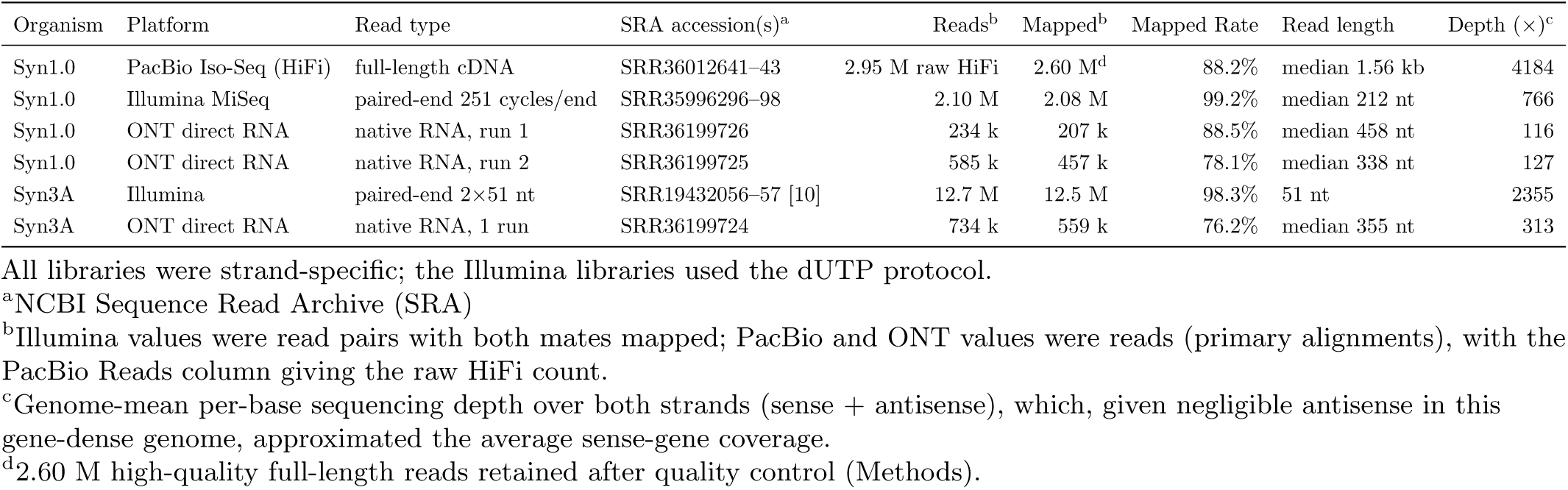
Summary of the RNA sequencing libraries for Syn1.0 and Syn3A.

| Organism | Platform | Read type | SRA accession(s)<sup>a</sup> | Reads<sup>b</sup> | Mapped<sup>b</sup> | Mapped Rate | Read length | Depth (×)<sup>c</sup> |
| --- | --- | --- | --- | --- | --- | --- | --- | --- |
| Syn1.0 | PacBio Iso-Seq (HiFi) | full-length cDNA | SRR36012641–43 | 2.95 M raw HiFi | 2.60 M<sup>d</sup> | 88.2% | median 1.56 kb | 4184 |
| Syn1.0 | Illumina MiSeq | paired-end 251 cycles/end | SRR35996296–98 | 2.10 M | 2.08 M | 99.2% | median 212 nt | 766 |
| Syn1.0 | ONT direct RNA | native RNA, run 1 | SRR36199726 | 234 k | 207 k | 88.5% | median 458 nt | 116 |
| Syn1.0 | ONT direct RNA | native RNA, run 2 | SRR36199725 | 585 k | 457 k | 78.1% | median 338 nt | 127 |
| Syn3A | Illumina | paired-end 2×51 nt | SRR19432056–57 [10] | 12.7 M | 12.5 M | 98.3% | 51 nt | 2355 |
| Syn3A | ONT direct RNA | native RNA, 1 run | SRR36199724 | 734 k | 559 k | 76.2% | median 355 nt | 313 |

All libraries were strand-specific; the Illumina libraries used the dUTP protocol.
<sup>a</sup>NCBI Sequence Read Archive (SRA)
<sup>b</sup>Illumina values were read pairs with both mates mapped; PacBio and ONT values were reads (primary alignments), with the PacBio Reads column giving the raw HiFi count.
<sup>c</sup>Genome-mean per-base sequencing depth over both strands (sense + antisense), which, given negligible antisense in this gene-dense genome, approximated the average sense-gene coverage.
<sup>d</sup>2.60 M high-quality full-length reads retained after quality control (Methods).

Two independent ONT direct-RNA runs of Syn1.0 were consistent with per-gene transcript abundances with Pearson *r* = 0.82 on log_10_. Per-gene transcript abundances were quantified using the gene-length corrected and normalized transcripts per million (TPM) metric (See Methods). However, they agreed only moderately in depth and read length in Table S2. This run-to-run variability of the ONT runs was expected from the different ribosomal-RNA depletion chemistries of the two libraries (Methods) and partly the short read length and the pervasive RNA processing described below.

Short-read Illumina is the standard for transcript quantification. The PacBio and Illumina TPMs were moderately consistent (Pearson *r* = 0.61, in line with previous reports [16]; Fig. S1b), with only a mild over-representation of long and low-abundance transcripts by PacBio (Fig. S1c,d). The two ONT runs correlated with Illumina to a similar degree. However, the correlation between two long-read methods, ONT and PacBio was weaker (Pearson *r ≈* 0.5). Illumina TPM was therefore used as the quantitative transcriptome axis for both Syn1.0 and Syn3A throughout, to avoid the bias introduced by different sequencing platforms. For Syn3A, only Illumina and ONT direct-RNA sequencing were available, with no PacBio library. The Syn3A ONT reads were short (median 355 nt, similar to the Syn1.0 ONT run 2 with the same protocol), too short to reconstruct full operon structures *de novo* as the PacBio isoforms did for Syn1.0, but sufficient to test the co-transcription of individual gene pairs, as applied in the genome-reduction analysis below. The Syn3A ONT and Illumina TPM agreed to a degree comparable to the cross-platform comparisons in Syn1.0 (Pearson *r* = 0.57; Fig. S1e).

### 459 operons in Syn1.0 annotated with full-length PacBio reads

PacBio reads were used to resolve the operons in the synthetic bacterium Syn1.0. After full-length non-chimeric (FLNC) recovery from PacBio cDNA reads and filtering, 2.6 M high-quality transcript reads gave an average sequencing depth of 4.2 k. To remove residual end noise, reads sharing transcript ends within 10 bp were clustered into 267 k isoform clusters, each a distinct transcript species with sharply defined boundaries (Methods).

Operons were segmented from these clusters in three stages (Methods). The 4 k clusters supported by at least 50 reads were grouped into 313 candidate operons by containment clustering [16] (See Methods). Adjacent same-strand operons that overlapped, or flanked a same-strand gene, were tested for read-level co-transcription and merged where continuous PacBio coverage bridged the junction, collapsing the set to 275. Finally, genes left uncovered by the resolved operons were rescued from spanning reads, the two highly structured rRNA operons were added directly, and 162 remaining isolated genes with low coverage were assigned as monocistronic operons. This produced a final map of 459 operons covering all 911 annotated Syn1.0 genes.

The resolved operons spanned a median 1,650 bp (mean 2,049 bp) and a mean of 2.0 sense genes (Fig. 1b,c). The largest operon was the *∼*11 kb ribosomal-protein supercluster *rpsJ* /0672*→secY* /0652, 20 ribosomal proteins plus SecY. In addition to the sense genes, 69 operons (15.0%) also enclosed a gene that was annotated on the opposite strand. Nine operons covered only antisense or intergenic regions, whose abnormal transcription activities are discussed later in Results.

Transcription start and termination sites (TSS, TTS) were read from the 127 canonical operons whose boundaries both fell in intergenic regions, where the ends reflected genuine start and termination positions rather than processed termini as shown in Fig 1a. Upstream of the TSS the bacterial *−*10 element was clearly recovered, with the TANAAT hexamer matching 87 of 127 operons (69%) and the extended TNNTANAAT motif 52 (41%), whereas the *−*35 region carried no fixed hexamer and was only broadly AT-rich (Fig. 1d). Signatures in both regions matched with those found in other bacteria [21, 22]. At the TTS, Rho-independent terminators [23] were found in 97 of 127 operons (76%). The canonical stem-loop architecture consisted of a stem-loop (median stem 8 bp, loop 4 nt) and the stems were enriched in G+C against the AT-rich genome (43% versus 24%) (Fig. 1d–f). The poly-U tract of median length of 6 was located immediately downstream of the stem-loop (Fig. 1g), and the mapped TTSs positioned a median of 1 nt (98% within 10 nt) beyond the poly-U tract. These conventional *−*10 and intrinsic-terminator elements indicated that the reduced genome retains a standard transcription-signal architecture, though the assignment was limited to operons with intergenic boundaries.

### Different sequencing methods disclosed complementary information on the division and cell wall cluster

Syn3.0 has pleomorphic morphology and may divide with primordial processes driven by only membrane curvature with a protein-lipid partition not being created [5, 6, 24, 25]. MMSYN1 0527, *ftsZ* /0522–*sepF* /0521– MMSYN1 0520, and other 15 genes were added to Syn3.0 to restore the normal morphology of Syn3A. We used all three sequencing methods to investigate the division and cell wall (*dcw* ) cluster *mraZ* /0525–MMSYN1 0520, together with its upstream co-transcribed genes *rpmF* /0526 and MMSYN1 0527, in Syn1.0 and Syn3A to support previous studies of the cell-division mechanism [5].

PacBio isoforms segmented the entire region into two operons, an upstream operon MMSYN1 0527– *rpmF* /0526–*mraZ* /0525–*mraW* /0524–*ftsA*/0523 and a downstream one *ftsZ* /0522–*sepF* /0521–MMSYN1 0520 in Fig 2. All Syn1.0 PacBio, ONT run1, and Illumina showed a jump of depth inside MMSYN1 0527, with transcription promoter -10 hexamer TATACT identified using PacBio 5*^′^* ends. Removing MMSYN1 0527 in Syn3.0 (now added back to Syn3A) would also remove this promoter and thus suppress the downstream genes, offering a transcription-level rationale for why MMSYN1 0527 had to be added back to build a minimal cell with normal morphology [5]. Of note, PacBio captured the shorter transcripts covering only *rpmF* /0526 less compared to ONT and Illumina, possibly due to the size selection of transcripts larger than 200 nucleotides in PacBio.

Cell division filament anchor gene *ftsA*/0523 had different coverage patterns. PacBio in Syn1.0 covered the first third with long reads starting from MMSYN1 0527 and the whole gene with shorter reads. ONT runs on Syn1.0 and Syn3A had transcripts with a median length of a few hundreds nucleotides and covered *ftsA*/0523 with consecutive reads. All libraries gave higher coverage on the cell division filament gene *ftsZ* /0522 than *ftsA*/0523 in both Syn1.0 and Syn3A, which was in line with tens more FtsZ proteins than FtsA from the proteomics.

Restoration of MMSYN1 0527 and the *ftsZ* /0522–*sepF* /0521–MMSYN1 0520 cluster in Syn3A recovered the transcription across the *dcw* cluster as shown in both the ONT and Illumina reads. Across *mraW* /0524– MMSYN1 0520, both the mRNA and protein fold changes between Syn1.0 and Syn3A stayed within 0.5 to 2.5 with a median near unity, indicating comparable expression in the two cells.

### Pervasive RNA processing truncates transcripts with a 3’ erosion bias in Syn1.0

We identified 459 operons in Syn1.0 that contained many isoforms shorter than the full operon length. When an isoform’s end falls inside the gene body, it most likely records post-transcriptional processing rather than an alternative promoter or terminator.

Two genes illustrated the opposite polarities of this truncation. In Fig. 3b, the neopullulanase gene MMSYN1 0178 exhibited substantial transcript erosion of the 3*^′^* end such that 49 of the 85 observed transcripts related to this operon displayed 3*^′^* processing, accounting for 29.5% of the reads covering this gene. Furthermore, unprocessed transcripts of this operon carried 65.5% and 4.6% only at the 5*^′^* end. In Fig. 3c, the leucyl-aminopeptidase gene *lap*/0154 whose transcription started 85 nt upstream with -10 box of TAAAAT, the polarity was reversed: 5*^′^* erosion of the 134 observed transcripts, accounting for 41.8% of the reads. Furthermore, unprocessed transcripts of this operon carried 49.7% of the operon’s total reads, and 8.2% only at the 3*^′^* end.

**Figure 3:**
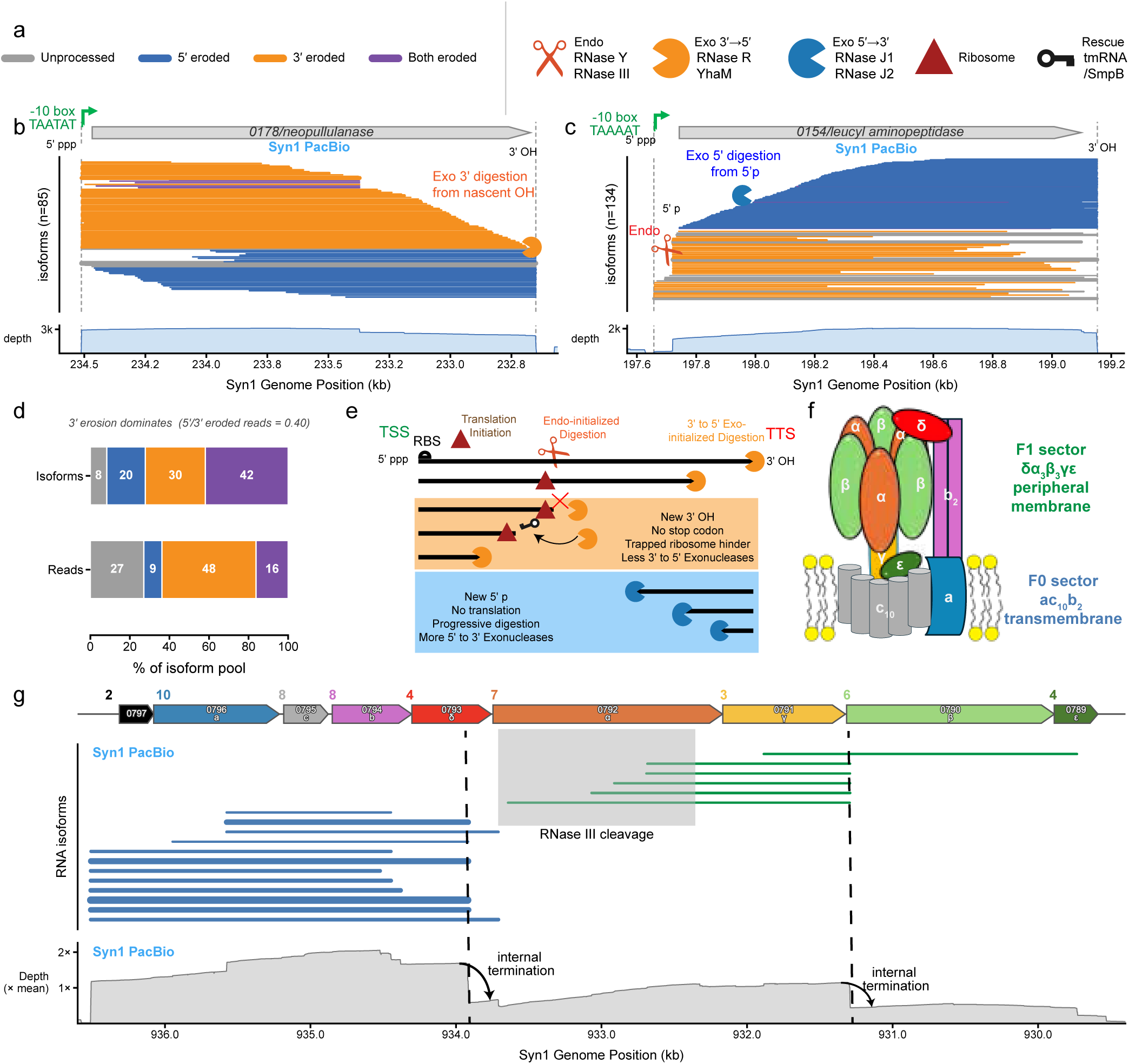
Pervasive and 3*^′^*-biased RNA processing reshapes the Syn1.0 transcriptome. (a) Shared legend for panel b to e: isoform end-context categories (left) and the ribonucleases, ribosome and trans-translation factors drawn in the schematic (right). (b) Isoforms at MMSYN1 0178 share the 5*^′^* start and erode at the 3*^′^* end (gene arrows, isoforms coloured by end-context, read depth). (c) Isoforms at *lap*/0154 share a 3*^′^* terminator and erode at the 5*^′^* end. (d) Transcriptome-wide composition of isoform end-context categories, by isoform kind and by isoform read; 3*^′^* erosion dominates. Isoforms were observed in PacBio. (e) Model of biased processing: endonucleolytic initiation (RNase III or RNase Y) followed by 3*^′^ →*5*^′^* exonucleolytic trimming, leaving stop-codon-less messages that are rescued by trans-translation; the 3*^′^ →*5*^′^* ribonucleases can also initiate RNA digestion directly at an accessible 3*^′^* end. RBS: ribosome binding site. (f) Subunit composition and spatial arrangement of the membrane complex ATP synthase in Syn1.0. (g) The ATP-synthase operon isoforms are organized as a 5*^′^* block (F_0_ plus *δ*) and a 3*^′^* block (F_1_ *γ*, *β*, *ɛ*) with clear separation in *atpA*/0792. Top 18 isoforms in this entire region were drawn with the line thickness scaling square root of the isoform reads. The digits (in the range of 2 to 10) before each gene denote the Shine-Dalgarno strength defined by the score of sequence alignment between the ribosome-binding sites and anti-Shine-Dalgarno sequence of the 16S rRNA.

This post-transcriptional processing was pervasive and strongly 3*^′^*-biased over the transcriptome in the PacBio dataset. Each of the 20,885 well-supported isoforms (*≥*10 reads) was classified by whether each end laid inside a coding sequence (a processing event) or in an intergenic region (a promoter or terminator). Only 8.3% of isoforms, carrying 27.0% of the reads, had both ends intergenic, thus classified as unprocessed primary transcripts (Fig. 3d). Among the processed majority, the asymmetry was pronounced: isoforms eroded only at the 3*^′^* end made up 30.3% of isoforms but 47.5% of reads, against 19.5% and 9.2% for those eroded only at the 5*^′^* end, so 5*^′^* erosion carried just 0.40 times the read weight of 3*^′^* erosion. As a result of the biased 3*^′^* truncation, 42.2% of the open reading frames contained within an isoform kept a start codon but had lost their stop codon.

The isoform endpoint asymmetry suggested the asymmetric activities of the resident ribonucleases. Syn1.0 retains a compact ribonuclease complement (Table S3): three endoribonucleases (RNase III, RNase Y, RNase M5), the 5*^′^ →*3*^′^* exoribonucleases RNase J1 and J2, and the 3*^′^ →*5*^′^* exoribonucleases RNase R and YhaM. Endonucleolytic cleavage by RNase III or RNase Y opens internal sites, and the fragments are trimmed exonucleolytically from their new ends [26]. Two asymmetries between the directions can account for the bias toward intragenic 3*^′^* ends as demonstrated in Fig. 3e. First, they initiate differently: RNase R can begin at an accessible 3*^′^* end such as a terminator poly-U tail in Fig. 3b, whereas RNase J1 requires a 5*^′^*-monophosphate end left by a prior cut and cannot initiate on an intact triphosphorylated 5*^′^* end in Fig. 3c. Second, 3*^′^* erosion in Syn1.0 was more limited than 5*^′^*. The processive enzymes differ in abundance: RNase J1 (92 copies, with RNase J2 as cofactor) is more than twofold more abundant than RNase R (36 copies, the only 3*^′^ →*5*^′^* enzyme that reads through structure, since YhaM stalls at a stem), so 5*^′^*-eroded fragments are cleared rapidly while 3*^′^*-eroded fragments persist and accumulate due to incomplete digestion. Furthermore, the action of 3*^′^* exoribonucleases could be hindered by stalled ribosomes, similar to how ribosomes protect 5*^′^* digestion intermediates [27]. The trans-translation system, tmRNA (*ssrA*/0158) and SmpB (*smpB* /0776), which rescues ribosomes stalled on the stop-codon-less messages 3*^′^* erosion products [28], is itself low in abundance (SmpB at 14 copies), so this rescue capacity may be limited.

**Table S3:** Ribonucleases and *trans*-Translation Rescue System in Syn1.

| LocusNum | Gene Product | Abundance | Function | Activity Signature |
| --- | --- | --- | --- | --- |
| *rnmV*/0003 | RNase M5 | 37 | Endo | 5S rRNA |
| *rny*/0359 | RNase Y | 70 | Endo | ssRNA w.<br>degenerate features |
| *rnc*/0418 | RNase III | 67 | Endo | dsRNA |
| *rnjA*/0600 | RNase J1 | 92 | Exo 5'→3' | 5' monophosphate |
| *rnjB*/0257 | RNase J2 | 142 | Exo 5'→3' | Cofactor of J1 |
| *rnr*/0775 | RNase R | 36 | Exo 3'→5' | ss 3' OH;<br>reads through structure |
| *yhaM*/0437 | YhaM | 117 | Exo 3'→5' | ss 3' OH;<br>stalls at structure |
| *ssrA*/0158 | tmRNA | - | Ribosome rescue | non-stop mRNA |
| *smpB*/0776 | Small protein B | 14 | tmRNA cofactor | non-stop mRNA |

RNA processing also remodeled co-expression within an operon. ATP synthase, a conserved membrane complex of subunit composition *α*_3_*β*_3_*γɛδac*_10_*b*_2_ [29], is organized into a transmembrane F_0_ sector (*ac*_10_*b*_2_) and a peripheral-membrane F_1_ sector (*δα*_3_*β*_3_*γɛ*) (Fig. 3f). The ATP-synthase operon (MMSYN1 0797– MMSYN1 0789) split into two non-overlapping isoform populations meeting at the *α*-subunit gene *atpA*/0792 (Fig. 3g). The upstream population covered the F_0_ subunits and *δ* (MMSYN1 0797–MMSYN1 0793) and ended at *atpA*/0792, while the downstream population began there and ran through the F_1_ subunits (MMSYN1 0791–MMSYN1 0798). Read depth fell from about 2*×* the genome-mean coverage of *∼*4,200 to roughly half across the boundary, indicating a possible internal termination event, though no intrinsic terminator was predicted. A similar drop recurred between *γ*and *β*. More interestingly, the isoforms spanning the *α* and *γ* subunits had varying 5*^′^* ends inside *atpA*/0792, the same gene whose ortholog in the Gram-positive *B. subtilis* carries two characterized RNase III cleavage sites [30].

### Transcript abundance is the strongest single determinant of the Syn1.0 proteome

Next, we sought to investigate the correlation between transcriptome and proteome in two synthetic organisms. Each protein’s intensity-based absolute quantification (iBAQ) was converted to a relative abundance in parts per million (iPM) and then to a copy number per cell (Methods). Of the 828 annotated proteins, 721 (87%) were detected. Copy number was strongly localization-dependent: cytoplasmic proteins had a median of 47 copies per cell (*n* = 516), against 21 for lipoproteins (*n* = 68), 10 for membrane proteins (*n* = 126), and 3 for extracellular proteins (*n* = 11), the most abundant being elongation factor Tu (*tuf* /0151) at roughly 7,200 copies (Fig. S2a). The low membrane and extracellular values likely reflected poor recovery of these classes by trypsin-only digestion rather than genuinely low abundance. Irrespective of subcellular localization, the 721 detected proteins’ abundances were significantly correlated with their respective transcript abundances quantified in Illumina short-read sequencing (Pearson *r* = 0.61, *R*^2^ = 0.38, *n* = 717). When considering only the cytosolic proteins, this correlation was strengthened to *r* = 0.70 (*R*^2^ = 0.49, *n* = 512; Fig. S2b).

**Figure S2:**
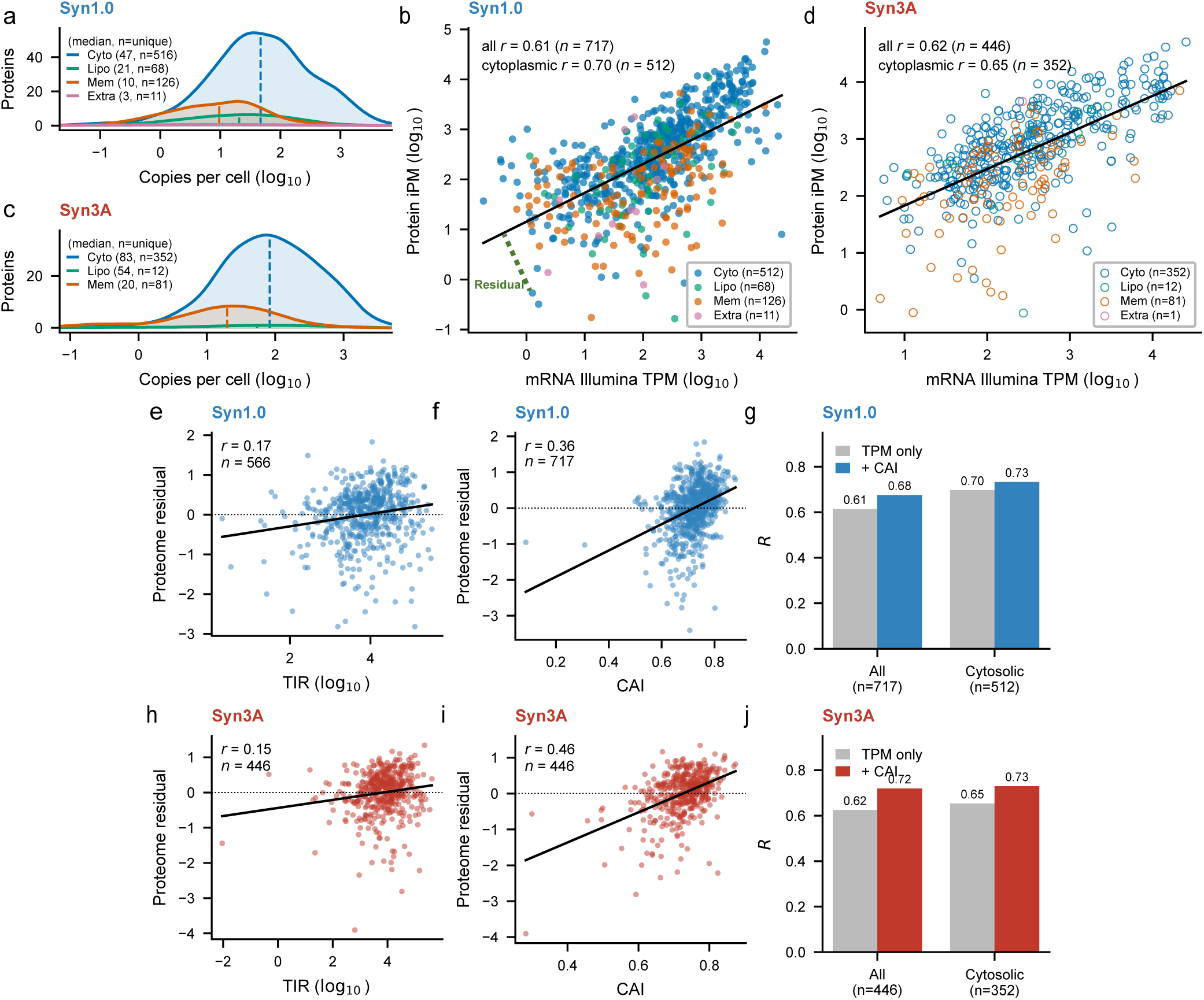
Supporting transcriptome–proteome analysis for Syn1.0 and Syn3A. Syn1.0 points are blue and Syn3A points red. The localization colours are shared between the two organisms. (a) Syn1.0 per-protein copy-number distribution by protein localization (Cyto, cytoplasmic; Lipo, lipoprotein; Mem, membrane; Extra, extracellular). (b) Syn1.0 mRNA Illumina TPM versus proteome iPM (log_10_), coloured by localization (filled circles). (c) Syn3A per-protein copy-number distribution by localization. (d) Syn3A mRNA Illumina [10] TPM versus proteome iPM (log_10_), open circles. (e) Syn1.0 predicted TIR (Translation Initiation Rate) versus the proteome residual. (f) Syn1.0 CAI (Codon Adaptation Index) versus the proteome residual. (g) Syn1.0 model Pearson *R* for the whole proteome and for cytoplasmic proteins, with and without CAI. (h) Syn3A predicted TIR versus the proteome residual. (i) Syn3A CAI versus the proteome residual. (j) Syn3A model Pearson *R* for the whole proteome and for cytoplasmic proteins, with and without CAI.

To dissect this residual, defined as the difference between protein’s log_10_ iPM and the value predicted from its linear transcript fit, two factors of translation initiation and translation elongation were tested. Translation-initiation rates (TIR) predicted with OSTIR [31] were largely uninformative, correlating only weakly with the residual (Pearson *r* = 0.17) and raising *R*^2^ by just 0.019, about 2% of the variance (Fig. S2e). This most likely reflected the limited accuracy of de novo initiation-rate prediction rather than an absence of initiation control as shown in another organism *Escherichia coli* [32]. Shine-Dalgarno strength, standby structure, RBS spacing and 5*^′^* mRNA folding, the four subfeatures constituting TIR all correlated poorly with the residual (Pearson *|r| ≤* 0.12; Methods). The codon adaptation index (CAI) [33], a proxy for elongation efficiency, was by contrast substantial: it correlated with the residual (Pearson *r* = 0.36; Fig. S2f) and increased *R*^2^ by 0.080, about 8% of the variance (Fig. S2g), so elongation efficiency approximated by codon usage against the AT-rich genome measurably tunes protein output.

Repeating the entire analysis in Syn3A returned the same picture (Fig. S2c-d,h-j). The one systematic difference was in absolute scale: although Syn1.0 carried about 1.27-fold more total protein molecules per cell (roughly 128,000 versus 100,000 in Syn3A, Methods), a comparable protein pool was distributed across fewer detected genes in Syn3A (446 versus 721), so its median per-protein copy number was nearly double that of Syn1.0 (66 versus 31 copies per cell in Fig. S2a,c). Transcript and protein levels correlated at Pearson *r* = 0.63 across the detected proteome (*n* = 446) and *r* = 0.65 for cytoplasmic proteins (*n* = 352), codon adaptation again explained a comparable slice of the residual (Δ*R*^2^ = +0.13), and predicted initiation stayed uninformative.

### Antisense and intergenic transcription in Syn1.0 trace largely to transcriptional noise and synthetic elements

Antisense transcripts are widespread in bacterial transcriptomes, but their functional significance remains uncertain. Lloréns-Rico *et al.* showed across 20 bacterial species and one chloroplast that antisense RNA abundance increases strongly with genomic AT content and concluded that most bacterial antisense transcripts arise from transcriptional noise initiated at spurious promoters, rather than representing regulated antisense RNA programs [37]. Their analysis included Syn1.0, providing an important precedent for interpreting low-abundance antisense transcription in this highly AT-rich genome.

Our full-length PacBio sequencing provided structural resolution of this phenomenon in Syn1.0. Antisense sequence was detected in only 1.4% of PacBio isoforms supported by *≥* 10 reads, which collapsed to 89 representative isoform clusters after merging transcripts with similar ends. These clusters were distinguished by the arrangement of sense and antisense segments (Fig. 4a): antisense-initiated transcripts from apparent spurious promoters (59/89, 66%), sense transcripts that read through an operon boundary into an antisense-annotated gene (26/89, 30%), and transcripts in which an antisense-annotated gene was embedded between two sense regions (4/89, 4%). The classes differed in length and abundance (Fig. 4b,c): embedded transcripts were longest (median 2.2 kb, versus 1.5—1.6 kb for the others) and read-through transcripts carried the highest typical support (median 111 reads). Every antisense cluster except one ( 31,700 reads in *his3* /) was far less abundant than the sense transcript it overlapped. These observations support the transcriptional-noise model proposed previously, while revealing the transcript architectures that generate the apparent antisense signal.

**Figure 4:**
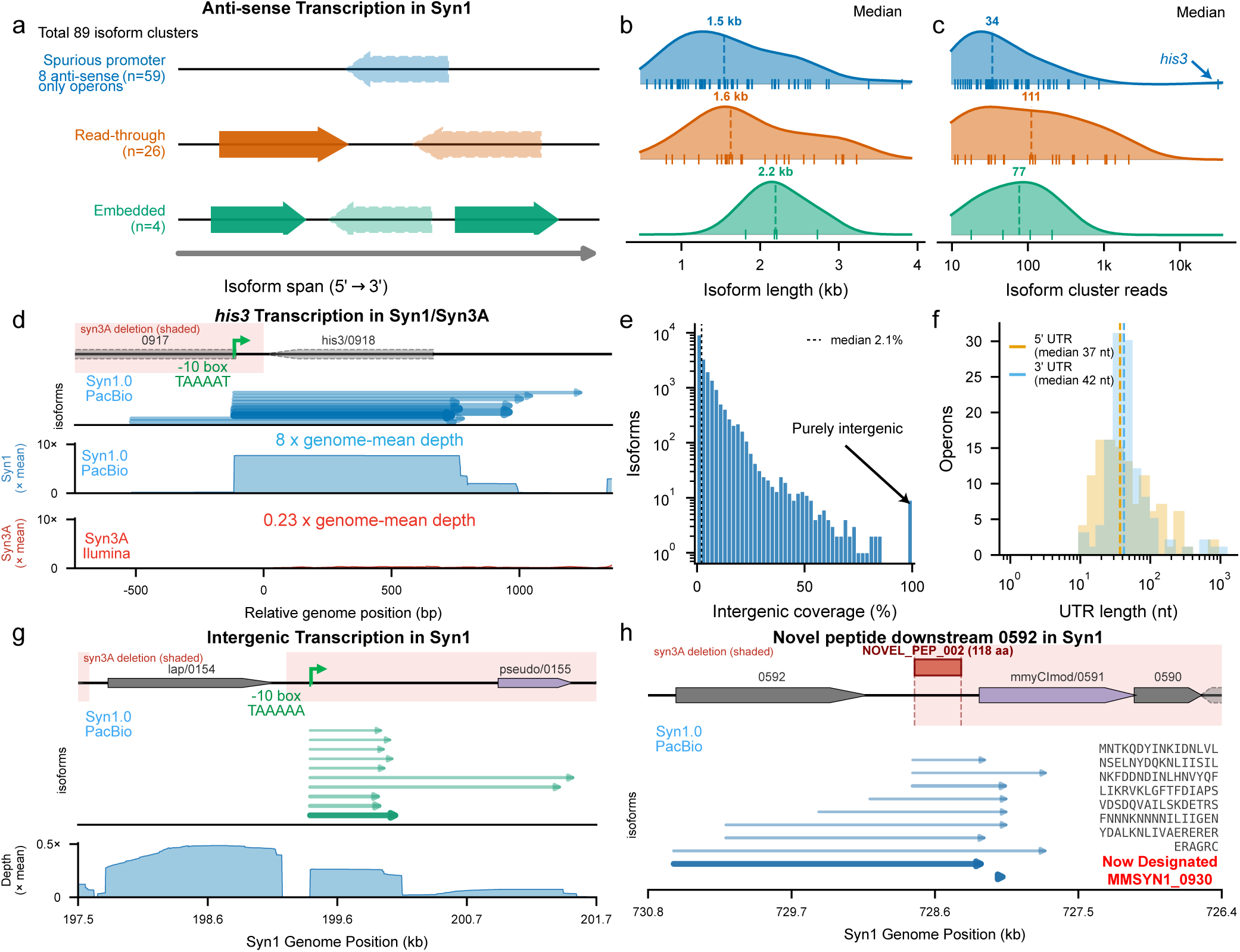
Non-canonical transcription and translation in Syn1.0. (a) The three classes of antisense transcription, spurious promoter, read-through, and embedded, relative to the sense isoform span. (b) Isoform length and (c) cluster-read support for each class, with the median marked. (d) Antisense transcription of the yeast vector gene *his3* /0918 in Syn1.0 and Syn3A: gene track, antisense isoforms in Syn1.0, and read depth of PacBio in Syn1.0 and Illumina in Syn3A [10], with the shaded band marking sequence deleted in Syn3A. Genome coordinates are relative to the 5’ end of *his3* /0918. Top ten abundant isoforms in this region drawn with the line thickness logarithmic scaling of the isoform reads. (e) Distribution of the intergenic coverage fraction across all isoforms. (f) 5*^′^* and 3*^′^* untranslated-region lengths of canonical operons. (g) An isolated intergenic transcript between *lap*/0154 and MMSYN1_0155. All ten isoforms with intergenic coverage larger than 50% in this region were drawn with line thickness logarithmic scaling of the isoform reads. (h) A novel 118-amino-acid ORF now designated as MMSYN1_930 near MMSYN1_0592 with its covering isoforms and peptide sequence. All ten isoforms with *≥* 10 reads that spanned the novel peptide were drawn with line thickness logarithmic scaling of the reads.

A particularly striking example is the yeast *his3* /0918 selection marker introduced during construction of the synthetic Syn1.0 genome as a artificial yeast gene [3]. This locus was transcribed almost entirely in the antisense direction, at a depth exceeding 30,000 reads and approximately eightfold the genome-wide average yet produced no detectable protein (Fig. 4d). The antisense transcripts originated near a TAAAAT sequence corresponding to a recognized *σ*-factor promoter motif. In Syn3A after genome minimization removed the upstream promoter-containing region, the *his3* coding sequence was retained, but the prominent antisense transcription was essentially lost, falling to 0.23-fold genome-average coverage. This provides a concrete example in which a synthetic genomic element generates abundant nonproductive transcription through a fortuitous promoter and in which genome minimization eliminates the transcription without eliminating the retained coding sequence.

Intergenic transcription was more pervasive than antisense transcription but was likewise generally low level. Across PacBio isoforms, the median fraction of sequence lying in intergenic regions was only 2.1%, arising predominantly from operon-flanking untranslated regions and short gaps within operons. Canonical operons had median 5*^′^* and 3*^′^* untranslated regions of approximately 37 and 41 nt, respectively (Fig. 4f). Only one long transcription unit was entirely intergenic: a cluster between MMSYN1 0154 and the pseudogene MMSYN1 0155 (Fig. 4g). Its absence of a canonical promoter *−*10 box and relatively low abundance were consistent with low-level transcription rather than a defined regulatory transcription unit.

To test whether abnormal antisense and intergenic transcription could encode proteins, every open reading frame within the 837 most abnormal isoforms, ranked by fraction of antisense and intergenic coverage, was scored for initiation with OSTIR [31]. This yielded *∼*29,000 candidate ORFs, of which the 100 highest-scoring candidates collapsed to 48 unique peptides (47 proteotypic). Re-examination of the Syn1.0 mass spectra confirmed two novel products: a 118-residue ORF downstream of MMSYN1 0592, now designated MMSYN1 0930 (Fig. 4h), and a 54-residue 5*^′^* extension of mmyCIVR/0768 (Fig. S3). The MMSYN1 0930 connects to the non-coding transcript MMYs 665 found with short-read Illumina sequencing by Lloréns-Rico *et al.* [37], and now with a novel peptide verified by proteomics. The second novel product represents an alternative protein form rather than a previously unannotated gene. Both novel peptides lie in restriction-modification loci adjacent to uncharacterized features and both occur in regions deleted in Syn3A.

**Figure S3:**
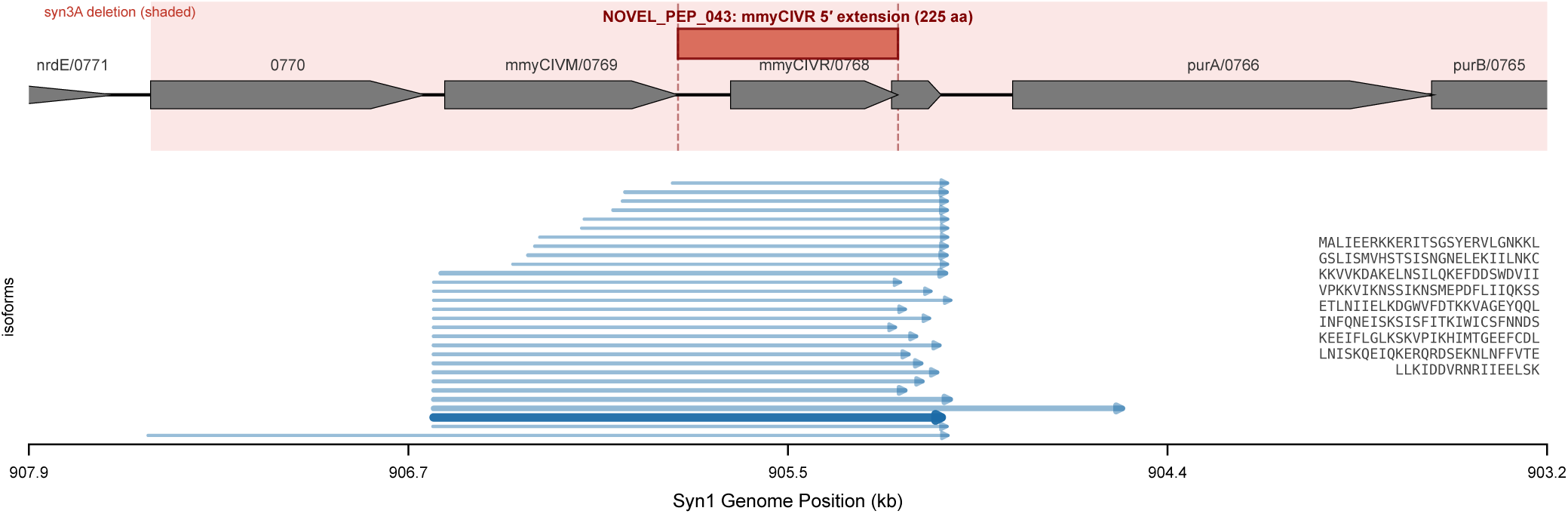
A new peptide found in Syn1.0 as 5*^′^* extension of *mmyCIVR*/0768. A novel protein product confirmed by mass spectrometry at *mmyCIVR*/0768: gene track (with the Syn3A-deleted region shaded), the PacBio isoforms fully covering the reading frame (*n* = 29, *≥*10 reads), and the 225-residue sequence. The *−* strand ORF (905,181–905,859) shares its stop codon with *mmyCIVR*/0768 but starts 162 bp (54 codons) further 5*^′^*, making it a 54-residue N-terminal extension of mmyCIVR; the entire *mmyCIV* cluster is deleted in Syn3A. All 29 isoforms with *≥* 10 reads that spanned the novel peptide were drawn with line thickness logarithmic scaling of the reads.

Together, these results indicate that the non-canonical transcription observed in Syn1.0 is dominated by low-level transcriptional noise, as predicted from the broader bacterial analysis [37], but that its sources can now be resolved at the level of individual full-length RNA molecules. In Syn1.0, these sources include fortuitous promoters in the AT-rich genome, transcriptional read-through, antisense annotations embedded within operons, and synthetic sequences introduced during genome construction. Genome minimization preferentially removed many of these transcriptional features, thereby simplifying the transcriptome in addition to reducing its coding capacity.

### Genome reduction disrupts the operons of key proteins

From Syn1.0 to Syn3A, the genome contracted from 1,078,809 to 543,379 bp through 95 deletions (*≥* 50 bp), removing *∼*536 kb while the retained backbone stayed 99.90% identical (Fig. 5a; Methods). Most deletions excised whole operons. Of the 459 Syn1.0 operons, over half lost every sense gene and a third kept all of theirs, leaving only *∼*50 partially truncated as leading, lagging or internal (Supplementary Data S3).

**Figure 5:**
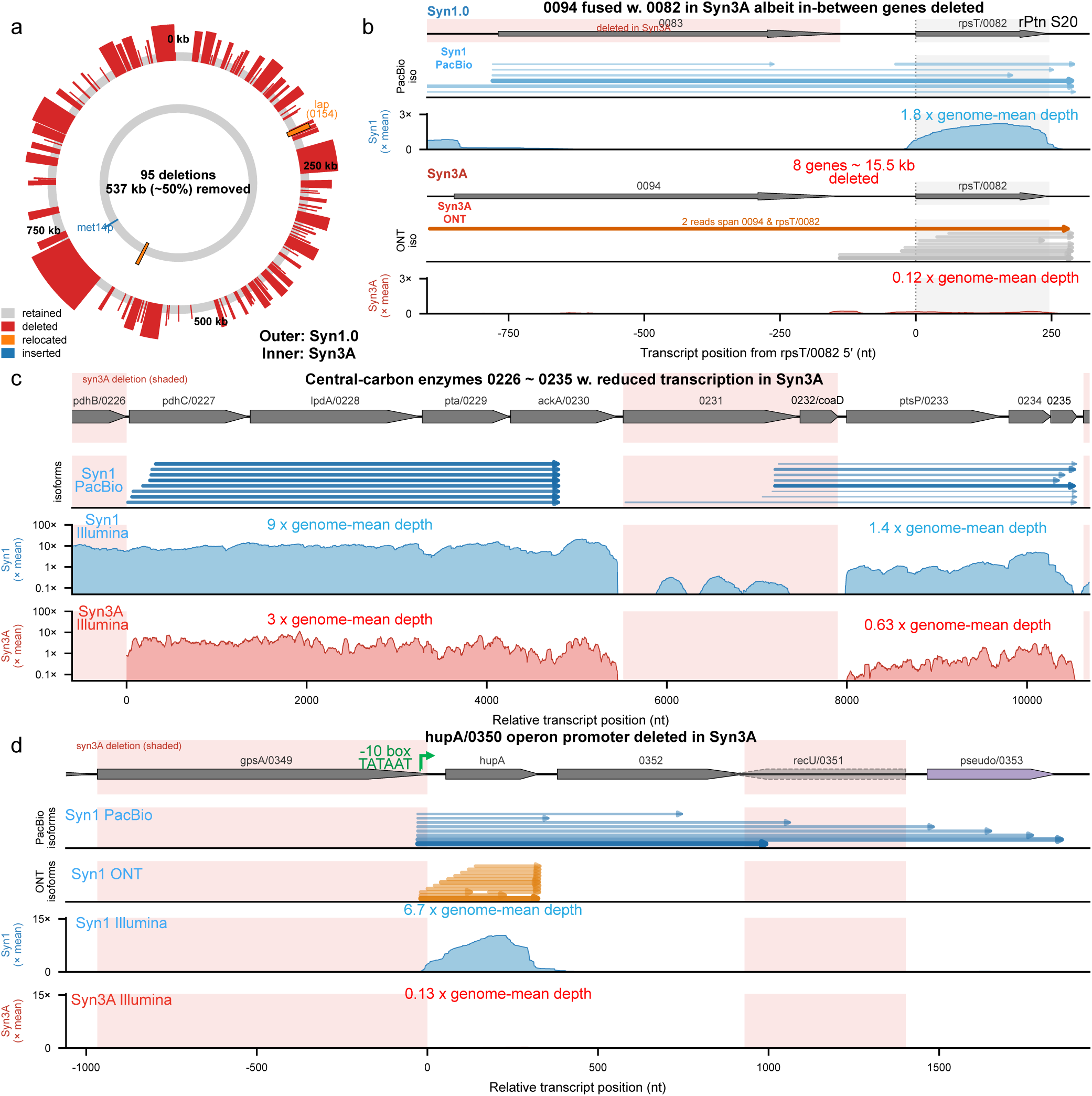
Genome minimization remodels Syn1.0 operon structure. (a) The Syn1.0 *→* Syn3A reduction: 95 deletions on the genome map. Each region is coloured by whether retained, deleted, relocated or inserted. (b) *rpsT* /0082 switched partner: co-transcribed with MMSYN1 0083 in Syn1.0, then fused downstream of MMSYN1 0094 in Syn3A after DEL 014 excises 15.5 kb (37 bridging ONT reads), both aligned on the shared *rpsT* /0082 5*^′^* end. Seven isoforms with reads *≥* in Syn1.0 PacBio and top eight abundant isoforms in Syn3A ONT were drawn. (c) Two disrupted central-carbon operons: deletions took the promoters of the pyruvate-dehydrogenase/acetate operon (*pdhC* /0227–*ackA*/0230) and the adjacent PTS operon (*ptsI* /0233–MMSYN1 0235), which drop together in Syn3A. Illumina data from Syn1.0 and Syn3A [10] were used for coverage. Top eight abundant isoforms in each operon were drawn with line thickness logarithmic scaling of the reads. (d) The *hupA*/0350 operon, disrupted by deletion of the promoter inside its neighbour MMSYN1 0349: PacBio isoforms fix the transcription start sharply within MMSYN1 0349, the Syn1.0 ONT 5*^′^* ends fall just downstream, and *hupA*/0350 collapses in Syn3A. Top eight abundant isoforms for Syn1.0 PacBio and 12 abundant isoforms for Syn1.0 ONT were drawn with line thickness logarithmic scaling of the reads. Illumina data of Syn3A [10] used.

Classifying every retained gene by how the reduction changed the nature of its transcription (see Methods) emphasized that only genes whose operon lost its own promoter with no replacement dropped robustly in expression (42 genes; median fold change 0.44 against 0.76 for structurally unaffected genes, two-sided Mann–Whitney *p* = 2.7 *×* 10*^−^*^4^), with no other class of contextual change differing significantly (Supplementary Data S3). Because the retained sequence is 99.90% identical between the two cells, this suppression is regulatory rather than mutational: deleting the promoter disrupts the operon and strands its retained genes without a transcription source.

Some proteins silenced this way are functionally essential. The nucleoid associated protein *hupA*/0350, Syn1.0’s principal chromosome-compacting factor, is transcribed from a promoter that lies inside its upstream neighbour MMSYN1 0349; the reduction deleted MMSYN1 0349 together with that promoter, disrupting the operon so that *hupA*/0350 collapsed in both transcript (fold change 0.020) and protein (0.014) (Fig. 5d). PacBio isoforms fixed this transcription start sharply. 79% of reads shared a single 5*^′^* base, from which a promoter with perfect TATAAT *−*10 box (consensus TANAAT) located just upstream yet removed in Syn3A. Syn1.0 ONT direct-RNA reads corroborated this start independently, their 5*^′^* ends falling within MM-SYN1 0349 as well but shifted roughly 15–30 nt downstream of the sharp PacBio start, possibly from 5*^′^* digestion of native transcripts.

Central-carbon metabolism was disrupted the same way at two adjacent operons. One deletion removed the promoter, along with the pyruvate-dehydrogenase E1 subunits *pdhA*/0225 and *pdhB* /0226, from the pyruvate-dehydrogenase/acetate operon. A second deletion removed the promoter of the neighbouring glucose-PTS operon, so the retained enzymes *pdhC* /0227–*ackA*/0230 and *ptsI* /0233–MMSYN1 0235 lost their promoters and fell together in Syn3A (Fig. 5c). This promoter loss is the transcriptional origin of the coordinated central-metabolism downshift quantified in the next section.

Aside from disrupting promoters, deletion events where two operons were fused into one across a deletion junction, were relatively rare (3 of 95 junctions; Methods) and ineffective at retaining transcriptional activity. The clearest case placed *rpsT* /0082 downstream of MMSYN1 0094 in a single unit (37 bridging ONT reads and 2 reads spanned both genes; Fig. 5b), though the in-between 8 genes were deleted. *rpsT* /0082 was transcribed only weakly in Syn3A (fold change 0.074) because the acquired MMSYN1 0094 promoter is itself weak. The similar case happened for another ribosomal-protein gene, *rpsO* /0294, thereby similarly reducing its transcription activity (fold change 0.144). Loss of a gene’s own promoter, by outright deletion or by replacement with a weaker fused one, rather than any change in the retained sequence, is thus the structural event that mainly suppresses some of Syn3A’s retained proteins.

### Genome minimization reallocates mRNA pool

Reducing the genome to half removed one fifth of the mRNA pool in Syn1.0. Of the 911 Syn1.0 loci, 418 are absent from Syn3A (382 protein-coding, 33 pseudogenes, 3 structural RNAs), together carrying 21.8% of the messenger-RNA pool and 22.3% of the proteome in Syn1.0 in Fig. 6a. Nearly half the genes thus held only about a fifth of the output, so the removed loci were well below average in expression, and their most transcribed members are all dispensable in rich medium: the yeast-vector marker *lacZ* /0915, the pyruvate-dehydrogenase subunits *pdhA*/0225 and *pdhB* /0226, and alanine dehydrogenase *ald* /0059 [2].

**Figure 6:**
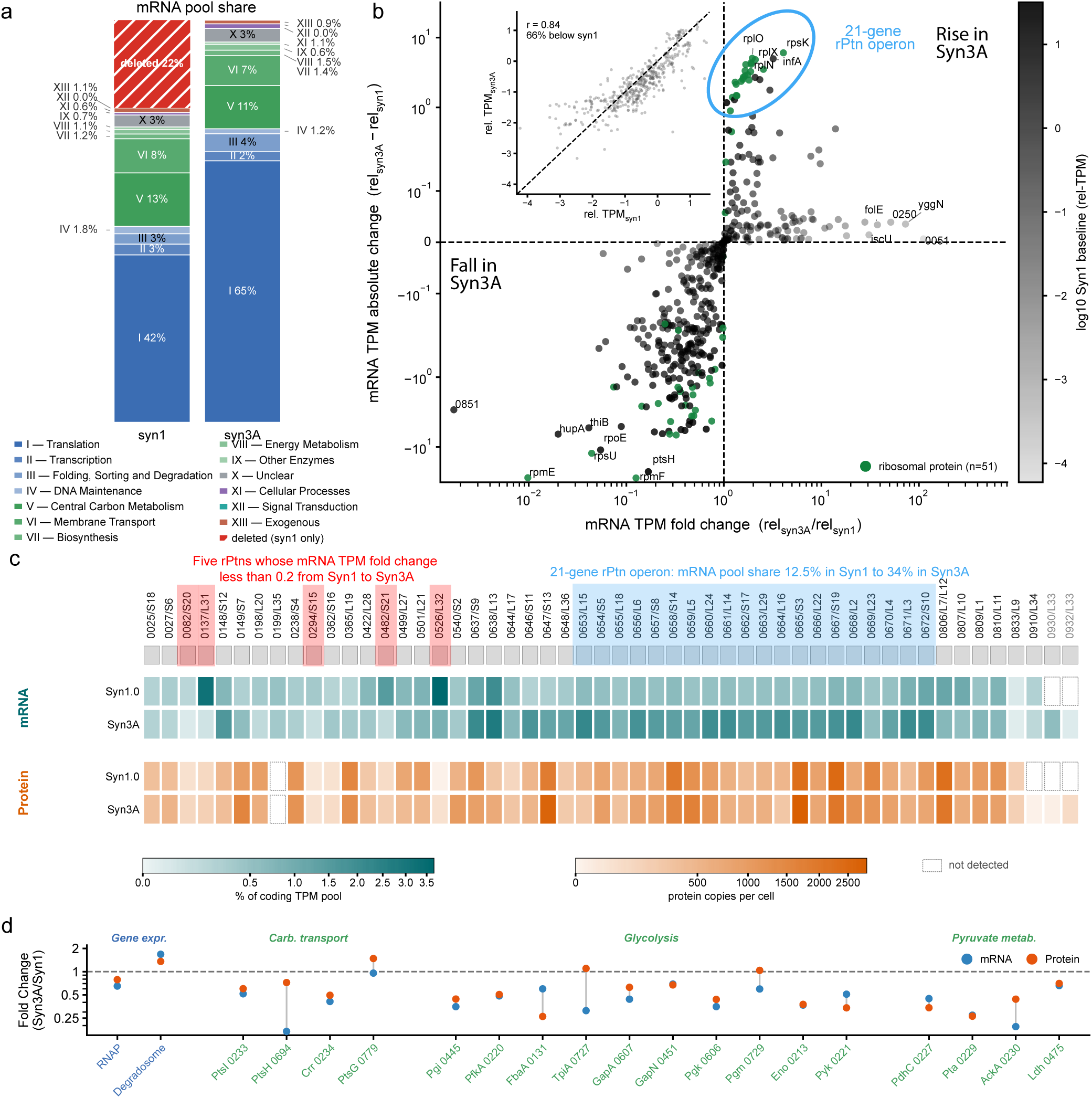
The Syn3A mRNA pool and proteome reallocate from Syn1.0 to Syn3A. (a) Composition of the messenger-RNA pool by Secondary function in Syn1.0 and Syn3A, quantified by Illumina. The category Translation includes ribosomal proteins, ribosome biogenesis, tRNA loading, tRNA biogenesis, and translation factors. Because the deleted genes contributed 21.8% of the Syn1.0coding-transcript pool, deletion alone would increase the fractional representation of the retained translation-associated genes from 42% to approximately 54%. The observed 64.9% in Syn3A therefore represents additional enrichment beyond the denominator effect of genome reduction. (b) Retained-pool mRNA fold change (Syn3A/Syn1.0) versus absolute change, the 51 ribosomal proteins highlighted; inset, the Syn1.0-vs-Syn3A relative-mRNA correlation (*r* = 0.84) in which two thirds of genes fall below the diagonal. (c) All 52 ribosomal-protein genes annotated in both cells. Top, gene panel labelled locus number/subunit-protein name; middle, messenger RNA as a share of each cell’s coding transcript pool; bottom, absolute protein copies per cell. Block opacity is proportional to the square root of the value. A dashed, unfilled block marks a gene’s mRNA or protein not detected. *rpmG* (JCVISYN3A_0930, JCVISYN3A_0932), the two extra L33 paralogues, are annotated only in Syn3A. Illumina of Syn1.0 (this study) and Syn3A [10]. (d) mRNA and protein fold changes of RNA polymerase, the degradosome, and the central-carbon enzymes. mRNA TPM quantified by Illumina.

The retained genes in Syn3A had similar transcript and protein abundances as in Syn1.0. Per-gene relative transcript levels in Syn1.0 and Syn3A correlated at Pearson *r* = 0.84 on log_10_ relative TPM (each gene’s TPM relative to the average gene TPM, which counteracts the different gene numbers of the two cells; *n* = 443 protein-coding genes in both organisms). The proteome repeated to a similar degree (*r* = 0.87 on log_10_ relative iPM, its protein analogue, *n* = 423). Comparisons are mean-normalized to the retained pool as explained in Methods.

A clear pattern of reallocation of mRNA pool was observed. The translation-associated genes occupied 64.9% of the mRNA pool now in Syn3A (Fig. 6a). The increase was driven predominantly by ribosomal-protein genes, including *rpsK* /0646, *rplO* /0653, *rplX* /0660 and *rplN* /0661, whose relative transcript abundances increased by 4.3 to 6.0 relative units. Most of the enrichment of ribosomal-protein genes was contributed by a single 21-gene ribosomal-protein operon, which accounted for 34% of Syn3A mRNA pool, compared to 12% in Syn1.0 in Fig. 6b,c. As the translation machinery occupied a larger fraction of the normalized mRNA pool, relative transcript abundance decreased for approximately two-thirds of the retained genes (see inset of Fig. 6b).

The *∼*11 kb ribosomal-protein operon MMSYN1 0652–MMSYN1 0672, a 21-gene polycistron from one promoter (a canonical TANAAT *−*10 box, TAGAAT) with no predicted internal terminator, now occupied one third of mRNA pool in Syn3A (Fig. S4). Full-length *∼*11 kb reads were rare at the read-length limit, but continuous coverage with no internal terminator marked it as one unit. The flanking deletions, which removed *dhaK* /0673 through MMSYN1 0677, brought a co-directional four-tRNA operon (MMSYN1 0678– MMSYN1 0681) to within 772 bp of its start, against 7,193 bp in Syn1.0, yet the operon’s own promoter was retained 179 bp from the nearest deletion. Despite this adjacency, the two were not co-transcribed (no Syn3A direct ONT RNA read spanned the junction; intervening short-read coverage 1.2% of the flanking depth; Fig. S4), so the operon’s dominance rested on its own retained promoter and the pool-level reallocation, not on read-through from the relocated tRNA genes. Despite the elevated transcript levels for this operon, mass spectrometry-based proteomics yielded similar ribosomal protein copy numbers.

**Figure S4:**
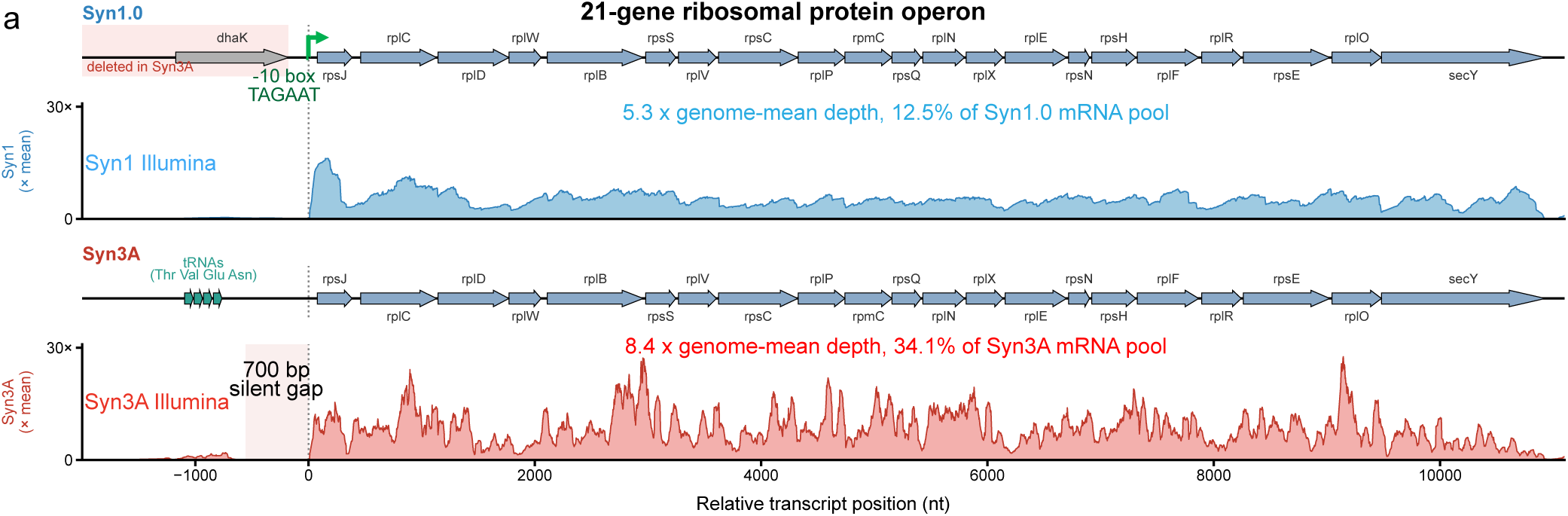
The 21-gene *∼*11 kb ribosomal-protein operon. The 21-gene *∼*11 kb ribosomal-protein operon (*rpsJ* /0672*→secY* /0652, minus strand) and its upstream neighbourhood in Syn1.0 and Syn3A, aligned on the shared operon 5*^′^* end (transcription start; gene tracks with Illumina depth, *×* genome-mean). Illumina data from Syn1.0 and Syn3A [10] was used.

In contrast to the long ribosomal-protein operon, several ribosomal-protein genes had reduced transcripts in Syn3A: the fusion-affected *rpsT* /0082 and *rpsO* /0294 instead collapsed (Fig. 5), and three structurally intact r-proteins (*rpmF* /0526, *rpmE* /0137, *rpsU* /0482) fell sharply in transcript despite unchanged context (Fig. 6b,c).

Two gene-expression machines moved in opposite directions, estimated from their limiting subunit (Fig. 6d). Core RNA polymerase (minimum of *rpoA*/2, *rpoC*, *rpoB* ) fell to a fold change of 0.65 in transcript and 0.79 in protein (35% and 21% reductions), whereas the RNase Y-based degradosome [38] (minimum of *rny*, *rnjA*, *yhaM* + *rnr* ) rose to 1.68 and 1.36 (68% and 36% increases). The reduced expression of transcription machinery together with increased RNA-turnover capacity is coherent with Syn3A’s longer cell cycle, roughly 105 min against 60 min for Syn1.0 [5].

In addition, central-carbon metabolism declined in concert at both mRNA and protein layers (Fig. 6d). Central Carbon Metabolism as a whole fell from 17.0% to 10.7% of the retained pool (median fold change 0.66, two-sided Mann–Whitney *p* = 1.3 *×* 10*^−^*^3^), with smaller losses in Membrane Transport (0.80, *p* = 0.016) and Transcription (0.55, *p* = 0.040). Among the glycolysis genes, *eno*/0213, *pgk* /0606, and *pyk* /0221 fell to transcript/protein fold changes of 0.37/0.38, 0.35/0.44, and 0.51/0.34; the pyruvate-dehydrogenase E1 subunits *pdhA*/0225 and *pdhB* /0226 were deleted outright while the retained E2 *pdhC* /0227 dropped to 0.45/0.34; and the acetate and lactate steps *pta*/0229, *ackA*/0230, and *ldh*/0475 fell to 0.27/0.26, 0.19/0.44, and 0.66/0.70. Part of this decline is the promoter-loss decapitation of the pyruvate-dehydrogenase and glucose-PTS operons shown earlier (Fig. 5), and the rest is the pool-level downshift onto which it is superimposed. This concerted downgrade predicts lower ATP and GTP output in Syn3A, a prediction awaiting an explicit metabolic-flux comparison.

## Discussion

Full-length sequencing resolved the transcriptome of Syn1.0 into operons and used it to analyze the transcriptome of the genetically minimized Syn3A. The transcriptomes were matched it to the proteomes in both Syn1.0 and Syn3A, showing that a synthetically-reduced genomes still runs a structured, heavily processed, and finely reallocated transcription pattern. We interpret the transcriptional organization based on the three questions raised at the end of the Introduction.

First, transcription is organized despite a low quantity of regulators encoded in the genome. Full-length PacBio isoforms defined 459 operons in Syn1.0, with matched promoter and terminator signatures for 127 operons (Fig. 1). Their transcripts were then pervasively reshaped by RNA processing, with markedly more 3*^′^* than 5*^′^* erosion. We read this asymmetry, tentatively, as the biased activities of exonucleolytic ribonucleases in Fig. 3. Discussion of the transcriptomics for Syn1.0 and Syn3A is further detailed based on the proteomics study in Fig. S2 (data in supplementary files). Furthermore, assessing the antisense and intergenic full-length transcripts against proteomics disclosed a new protein-coding gene, MMSYN1 0930, that was previously identified as non-coding region [37] in Fig.4(h).

Second, genome minimization from Syn1.0 to Syn3A was structurally conservative yet functionally transformative for the retained genes. The minimization preserved gene order and largely excised whole operons, and some deletions we observed include antisense and intergenic transcripts that we traced to inherited *M. mycoides* mis-annotation and synthetic-construction artifacts in Fig. 4. Where deletions fell inside a retained operon they occasionally removed a gene’s own promoter and disrupted its transcription, the clearest case being the nucleoid associated protein HupA, whose promoter lies within the deleted *gpsA*/0349 and whose transcript abundance accordingly collapses in Syn3A in Fig 5. The deletion pattern back in 2016 [4] was based on gene coding regions without knowing the exact promoters and terminators that we found with long-read sequencing in this study. As such, MMSYN1 0527 was deleted in Syn3.0 [4] but had to be added back to Syn3A [5] to restore the transcription of its downstream *dcw* cluster genes (Fig. 2). Had long-read sequencing information been available, different minimization decisions might have been made.

Beyond these losses, the retained transcriptome was reallocated toward the single 11 kb ribosomal-protein operon, which tripled its share of the mRNA pool, while RNA polymerase and the central-metabolic enzymes fell and the RNA degradosome rose in Fig 6. Of note, we identified the particular example of fructose-1,6-bisphosphate aldolase encoded by *fbaA*/0131, whose protein abundance had to be manually increased to allow greater glycolysis flux in the forward direction when parameterizing the glycolysis in the WCM of Syn3A [12]. Two non-exclusive effects could drive the 11 kb ribosomal-protein operon’s transcript abundances, a tRNA operon that minimization brought adjacent to it raising the local availability of RNA polymerase without read-through, and the loss of HupA [8] shifting genome-wide supercoiling, making the search for strong promoters, like the ribosomal-protein operon one, easier.

Third, these expression changes point to molecular origins for the phenotypic differences between the two cells. Ribosomal proteins S15 and S20 as primary binders to 16S rRNA [39, 40] have largely reduced mRNA abudance in Syn3A in Fig. 6c, thus preventing the assembly of ribosome small subunit at the nucleation stage. Transcription of ribosomal proteins L31 and L32 also collapsed in Syn3A, yet these two proteins came late in the assembly of ribosome large subunit [41, 29, 6]. A preliminary study on roughly 250 tomograms [42, 43] of Syn3A revealed that 100 ribosome large subunits were free with 170 complete ribosomes per tomogram, which limited the protein synthesis capacity in Syn3A. The coupling of lower transcription and energy output to faster mRNA turnover by the degradosomes in Fig.6 is also consistent with Syn3A’s roughly twofold longer cell cycle [2, 5]. And the abundance collapse of the nucleoid associated protein, HupA noted above may account for Syn3A lacking the persistent chromosome supercoiling [8].

All three RNA sequencing techniques—PacBio, ONT and Illumina—were used together to answer these questions. PacBio cDNA Iso-seq gave the highest coverage and the longest isoforms, from which the co-transcription operons were annotated. ONT direct-RNA libraries provided complementary yet slightly different information. In the *dcw* cluster (Fig. 2), ONT and Illumina correctly sampled much of the short *rpmF* /0526 transcripts, whereas PacBio failed possibly due to the size selection in the Iso-seq library preparation. Both long-read sequencing revealed that the promoter for *rpmF* /0526 is located within its upstream 0527 in Fig. 2 and the promoter for *hupA*/0350 is located within its upstream *gpsA*/0349 in Fig. 5(d). However, direct ONT libraries gave shorter reads than PacBio cDNA Iso-Seq, plausibly because the same pervasive processing degrades these native transcripts, so degradation-prone messages would benefit from a better RNA-protection protocol. To compare the Syn1.0 and Syn3A transcriptome, quantifications were done using the same Illumina short-read platform to reduce the bias introduced by different sequencing. Several limitations remain. We did not have PacBio data for Syn3A, so its operon-level changes are inferred from shorter ONT reads and await full-length confirmation. Effective rRNA depletions were used in all sequencing library preparations, and fraction of rRNAs lower than 6% were observed for all (Methods). Thus, expression here was framed as a share of the mRNA pool and excluded rRNA. Mass spectrometry-based proteomics were reported to be less accurate on the ribosomal-protein abundances [44, 45]. Going from Syn1.0 to Syn3A, the ribosomal proteins encoded by the giant 21-gene operon maintained similar copy numbers from the proteomics data, yet their share of the mRNA pool more than doubled. Ribosome profiling [46, 32] can sharpen those protein abundances and probe how their translation profiles change as a result of the mRNA pool reallocation.

More broadly, the reallocation of resources between transcription and translation echoes the balancing of cellular economy seen when other genomes are streamlined, such as *B. subtilis* [47], suggesting a general principle rather than a Syn1.0 and Syn3A idiosyncrasy. The isoform-resolved transcription in ATP synthase operon echos that differential transcription, RNA stability, and translation coherently contributed to the stoichiometric protein synthesis as in *B. subtilis* [48]. The analysis and visualization of transcription in all genomic regions in Syn1.0 and Syn3A are shared via Jupyter Notebook. Together, this operon-resolved map of gene co-expression and transcript quantification supplies the RNA-level foundation that a four-dimensional whole-cell model of the minimal cell has so far lacked [6].

## Methods

### Comparison of PacBio, ONT, and Illumina RNA sequencing techniques

Three complementary RNA sequencing (RNAseq) platforms were used to profile the transcriptome of Syn1.0 and Syn3A, each capturing a different facet of transcript structure and abundance. Illumina short-read sequencing provided the quantitative backbone, PacBio Iso-Seq^TM^ resolved full-length isoform structure, and Oxford Nanopore Technologies (ONT) direct-RNA sequencing provided native, amplification-free long reads.

Illumina-based sequencing is currently the most common technique, owing to its robustness, cost, and throughput. Illumina instruments generate billions of reads per flow cell, and the chemistry has been refined considerably since the technique was introduced [49]. Library preparation requires several intermediary steps prior to sequencing: (i) fragmentation of RNA, (ii) conversion of isolated mRNA fragments to double-stranded complementary DNA (cDNA) by reverse transcription, (iii) adapter ligation, (iv) PCR amplification, and (v) size selection [15]. Fragments are sequenced from one end (single-end) or both ends (paired-end), typically with read lengths of tens of nucleotides depending on the instrument. The reads are short but exceed 99% base accuracy, particularly with the latest chemistries and instrumentation.

Pacific Biosciences (PacBio) sequencing requires similar library-preparation steps from mRNA to cDNA but sequences full-length cDNA through the Iso-Seq^TM^ protocol, which incorporates SMRTbell adapters for circular consensus sequencing (CCS, also known as HiFi sequencing) [50]. Typically, the process of removing short fragments were used in library preparation to maximize the read length and data yield. Improved instrumentation such as the PacBio Revio and concatemerization techniques such as Kinnex yield accurate full-length isoforms of up to 20 kb, although longer transcripts may be truncated or lost when size selection is applied. PacBio requires substantial RNA input, so PCR amplification is commonly used during library preparation, and the platform carries a known bias toward smaller fragments that can complicate expression quantification [51].

Both Illumina and PacBio Iso-Seq convert RNA to cDNA and rely on PCR amplification and size selection, and short reads alone cannot capture the full complexity of the transcripts present in a bacterial cell. ONT instead captures long transcripts directly by passing native RNA through a membrane-embedded nanopore (3*^′^* to 5*^′^*) and reading the perturbations of an applied electric current [52]. Direct-RNA sequencing was included for its ability to read full-length native transcripts without conversion to cDNA, which can perturb the native molecule and lose information [53]. Beyond quantification, native long reads resolve gene co-expression, antisense and intergenic transcription, and, in principle, common base modifications such as N6-methyladenosine [54].

The comparison therefore reveals an important tradeoff between transcript structure and transcript population representation. PacBio Iso-Seq provided highly accurate long-read information for transcripts represented in the cDNA library, whereas ONT direct RNA sequencing recovered native RNA populations that were poorly represented in the cDNA-derived datasets. Importantly, ONT direct RNA sequencing should not be assumed to provide quantitatively more accurate transcript abundance measurements than PacBio or Illumina. Previous systematic comparisons in *Escherichia coli* showed that ONT cDNA-based protocols can provide greater read yield and per-read accuracy than ONT direct RNA sequencing, emphasizing that direct RNA sequencing has important technical limitations of its own [17].

### Illumina Short-read Sequencing of Syn1 Transcriptome

#### Illumina RNA Sample Preparation

Sample preparation was conducted following [55] A frozen stock of Syn1.0 strain was provided by the JCVI. The frozen cell stock was scraped and inoculated in 10 mL of SP4+KnockOut medium under BSL2 conditions and incubated in a horizontal rotating shaker at 37°C for about 24 hours until the culture medium turned to orange. The SP4 media is made from a two-part formulation: part ***I*** –3.5 g Mycoplasma Broth Base, 10 g Bacto Tryptone, 5.3 g Bacto Peptone, and 600 mL distilled water mixed to suspend well, adjusted to pH 7.5, and then autoclaved 15 min at 121°C and part ***II*** –25 mL Glucose 20% w/v stock, 50 mL CMRL 1066 (10X stock w/o phenol red, w/o bicarb, w/o Gln), 14.6 mL sodium bicarbonate 7.5% w/v stock, 5 mL L-glutamine 200 mM stock, 35 mL Yeast extract solution, 100 mL TC Yeastolate 2% w/v stock, 170 mL Serum (heat inactivated FBS, HS) or substitute (KO), 2.5 ml Penicillin G (400,000 U/mL stock), and 1.5 mL Phenol red (1% w/v) filtered and sterilized (0.2 *µ*m) then stored at 4°C. To complete the SP4 medium, combine 1.5 volume of part ***I*** with 1 volume of part ***II*** .

RNA purification and depletion steps were conducted using sterile, RNase-free pipette tips (Thomas Scientific 1145N16, VWR 76322-134) and microcentrifuge tubes (Posi-Click 1149K01) on a countertop or biosafety cabinet treated with RNase decontamination solution (Invitrogen AM9780). Solutions apart from those provided with the kits were prepared using DEPC-treated water (Invitrogen 46-2224). RNA was extracted from JCVI-syn1.0 cells using the PureLink RNA Mini Kit (Invitrogen 12183018A) according to the manufacturer’s protocol. Briefly, *≤* 1 *×* 10^9^ cells (2 mL of exponential phase culture) were harvested (4000*×*g, 5 min, 4°C) and resuspended in lysozyme solution (100 *µ*L; 10 mM Tris-HCl, pH 8.0, 0.1 mM EDTA, and 20 ^mg^⁄_mL_ lysozyme Egg white (L-040-25, Goldbio). SDS solution (500 *µ*L; 10% w/v) was added and the cells were incubated for 5 min at room temperature (rt). Lysis buffer containing *β*-mercaptoethanol (350 *µ*L) was added and the sample was vortexed and passed six times through an 18-gauge needle. The resulting homogenate was centrifuged (12,000*×*g, 2 min, rt) and 100% ethanol (250 *µ*L) was mixed with supernatant separated from the precipitate. The sample was purified through a spin cartridge provided with the kit and eluted with 2*×*100 *µ*L DEPC-treated water. To remove DNA, 1/9 volume of 10X DNase I Buffer (NEB, B0303S) and DNase I (NEB M0303S; 4 U total) were added and the mixture was incubated for 10 min at rt. EDTA (5 mM final concentration) was then added, and the reaction was heat-inactivated for 10 min at 75°C. Quantification with a NanoDrop Lite Spectrophotometer (Thermo Fisher ND-LITE-PR) typically showed yields of 40-60 *µ*g of RNA.

Ribosomal RNA was removed from the DNase-treated RNA samples using the RiboMinus Bacteria 2.0 Transcriptome Isolation Kit (Invitrogen A47335) according to the manufacturer’s protocol. Briefly, *≤*5 *µ*g of DNase-treated RNA was mixed with 2X Hybridization Buffer (50 *µ*L) and RiboMinus Pan-Prokaryote Probe Mix (3 *µ*L) to a final volume of 100 *µ*L. The RNA/probe mix was denatured for 10 min at 70°C, followed by hybridization of the rRNA and probes for 20 min at 37°C. The RNA/probe mix was added to a microcentrifuge tube containing a 200 *µ*L suspension of RiboMinus Magnetic Beads in 1X Hybridization Buffer and incubated for 15 min at 37°C. The tube was placed on DynaMag-2 Magnet stand (Thermo Fisher 12321D) for 1 min. The supernatant (300 *µ*L) containing the rRNA-depleted RNA was carefully aspirated and mixed with Nucleic Acid Binding Beads (10 *µ*L), Binding Solution Concentrate (400 *µ*L), and 100% ethanol (1 mL). The mixture was incubated for 5 min at rt and placed on the magnetic stand for 5 min. The supernatant was discarded, the beads were washed with Wash Solution (300 *µ*L) and placed on the magnetic stand again. The supernatant was discarded, the beads were air-dried for 5 min, and the RNA was eluted with DEPC-treated water heated to 70°C. The Nucleic Acid Binding Beads were removed using the magnetic stand, and the supernatant was transferred to a fresh microcentrifuge tube.

#### Illumina RNA Sample Quality Control Analysis

Quantification of RNA was conducted using the Qubit RNA High Sensitivity assay (Invitrogen Q32852) and measured on a Qubit fluorometer (Invitrogen). Within-range samples treated with the RiboMinus kit were monitored with an AATI 5200 Fragment Analyzer (Agilent) showing depletion of rRNA-associated peaks. Method name: DNF-472T22 - HS Total RNA 15nt.mthds; Gel prime: No; Full conditioning: Yes; Gel prime to buffer: Yes; Gel selection: Gel 2; Perform prerun: 7.0 kV, 30 sec.; Rinse: No; Marker 1: No; Rinse: Tray: 3, Row: A, Dip count: 2; Sample injection: 6.0 kV, 150 sec.; Separation: 7.0 kV, 31.0 min.; Tray name: Tray-3; Analysis mode: RNA (Eukaryotic).Quality control plots of RNA libraries have been included in Supplementary Data S4.

#### Illumina Library Preparation and Sequencing

The RNAseq libraries were prepared with the Kapa Hyper Stranded mRNA library kit (Roche). The library pool was quantitated by qPCR and sequenced on one MiSeq Nano flowcell for 251 cycles from each end of the fragments using a MiSeq 500-cycle sequencing kit version 2.

#### Illumina MiSeq Read Processing, Mapping to the Genome

Three strand-specific paired-end RNA-seq datasets of Syn1.0 were retrieved from the NCBI SRA database (accessions SRR35996296, SRR35996297, and SRR35996298), sequenced on an Illumina MiSeq for 251 cycles from each end as described above. Datasets SRR35996296 and SRR35996297 are technical replicates of one biological RNA sample, and SRR35996298 is a second biological sample, yielding *∼*505 k, *∼*511 k, and *∼*1.09 M read pairs, respectively. The deposited reads are of variable length: they were demultiplexed with bcl2fastq, which emits 250 nt reads and trims the 3*^′^* adapter at demultiplexing, and because the library inserts (mean 214–234 nt across the three datasets) are shorter than the read, most reads are clipped back to the insert length. Pooled over the three datasets the reads span 35–250 nt (median 212 nt, mean 198 nt), with 22.4% at the full 250 nt and a floor at 35 nt set by the bcl2fastq minimum trimmed-read length.

Deposited reads were aligned to the Syn1.0 reference genome (NCBI accession CP002027.1) with bowtie2 v2.5.5 [56] in default paired-end mode, against an index built with bowtie2-build from the same reference. Overall alignment rates were 99.49–99.56% across the three datasets, of which 89.4–93.1% of pairs aligned concordantly exactly once and 5.1–7.8% aligned concordantly more than once. SAM output was converted, sorted, and indexed with samtools v1.23 [57]. Strand-specific BAMs were generated using the dUTP / fr-firststrand flag convention (read 2 forward on the + strand and read 1 reverse on the + strand define plus-strand reads; their reverse complements define minus-strand reads), matching the Kapa Hyper Stranded mRNA library preparation, and per-strand sequencing-depth bedGraph tracks were computed with samtools depth. Between 3% to 6% of aligned bases fell within the two rRNA operons across the three Syn1.0 Illumina datasets, indicating effective rRNA depletion.

#### Quantification of TPM from Sequencing Depth

Per-gene transcript abundances were quantified from the strand-specific depth tracks rather than from read counts. For each gene annotated in the Syn1.0 reference (feature type gene), a sense and an antisense mean per-base depth were computed over the gene body from the plus- and minus-strand bedGraph tracks using prefix sums, with the sense strand taken as the gene’s annotated strand. Dividing the summed depth by the gene length, 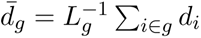, length-normalizes coverage, and each value was scaled to transcripts per million as 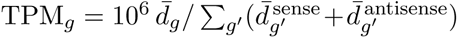 using a single sense-plus-antisense denominator per dataset.

TPM was computed independently for the three datasets, which agreed well (Pearson *r* = 0.98 between the two technical replicates and *r* = 0.92–0.93 between biological samples). All TPM correlations reported here are Pearson correlations of log_10_ TPM over the genes with non-zero TPM in both libraries compared. The datasets were merged by a two-step average, first across the technical replicates within their shared biological sample and then across the two biological samples at equal weight, yielding a representative Syn1.0 Illumina TPM profile (available in Supplementary Data S2).

### PacBio Long-read Sequencing of Syn1 Transcriptome

#### PacBio RNA Sample Preparation

Sample preparation was conducted in the same manner as in Illumina, except DNase-I treated RNA was purified by spin cartridge (GeneJET RNA Purification Kit, Fisher K0731) instead of heat-inactivated with EDTA. In addition, prior to rRNA depletion, DNase I-treated RNA was polyadenylated using 1-10 *µ*g of the RNA, ATP (1 mM final concentration) (NEB B0756), *E. coli* Poly(A) Polymerase (NEB M0276S), and *E. coli* Poly(A) Polymerase Reaction Buffer (B0276S). The reactions incubated at 37°C for 20 minutes and then purified using the GeneJET RNA spin cartridge.

#### PacBio Library Preparation and Sequencing

Ribodepleted and polyadenylated RNAs were purified using an RNA Clean & Concentrator-5 column (Zymo Research, Cat. #R1015) following the protocol designed to remove fragments smaller than 200 nucleotides. Twenty nanograms of purified RNA were converted into complementary DNA (cDNA) using the IsoSeq Express Oligo Kit from Pacific Biosciences. The cDNA molecules were barcoded with the Barcoded Overhang Adaptor Kit 8A and further processed into a library using the SMRTbell Prep Kit 3.0 (Pacific Biosciences). The resulting libraries were pooled at equimolar concentrations, and the pool was sequenced on one SMRT Cell 8M using a PacBio Sequel IIe system in CCS sequencing mode with a 30-hour movie.

#### PacBio RNA Sample Quality Control Analysis

RNA-sample QC was conducted in the same manner as in Illumina, and the resulting cDNA and final libraries were additionally sized on an Agilent Femto Pulse. Quality control plots of RNA libraries have been included in Supplementary Data S4.

#### PacBio Iso-Seq Read Processing, Mapping to the Genome, and Quality Control

Three technical replicates were generated for one biological sample of the synthetic bacterium JCVI-Syn1.0 (NCBI SRA accessions SRR36012641, SRR36012642, and SRR36012643). The three replicates were pooled into one read file of 2.95 M entries given their high consistency. Demultiplexed PacBio HiFi cDNA reads were preprocessed to get full-length non-chimeric (FLNC) reads with a custom streaming pipeline before isoform clustering. The pipeline comprises four sequential steps: strand orientation, primer removal, 3*^′^* poly(A) trimming, and post-mapping quality control. Each step operates on FASTQ file; no third-party Iso-Seq tool (e.g. lima, isoseq3 refine) was used, giving explicit control over filter thresholds in this bacterial, non-natively-polyadenylated context.

Each double-stranded cDNA read carries one of two primer configurations: an antisense molecule reads 5*^′^*–H1 (oligo-dT handle, TAAGCAGTGGTATCAACGCAGAGTAC, 26 nt)–polyT–[reverse-complement of mRNA]–BCRC–3*^′^*, whereas a sense molecule reads 5*^′^*–BC (barcode, CCCAACCCTGCGACTTCATTGCA, 23 nt)–mRNA–polyA–H2–3*^′^*. A read was reverse-complemented to sense orientation if H1 occurred within its first 80 bp or if it ended in BCRC; otherwise it was kept as-is. Oriented reads were then required to terminate in an exact polyA (*≥* 10 nt) + H2 suffix as evidence of a full-length sense cDNA. Of 2.95 M raw HiFi reads, 1.35 M were flipped via H1 and 191 k via BCRC; 255 k lacking the canonical polyA+H2 suffix were discarded, yielding 2.69 M oriented reads.

H2 was trimmed from the 3*^′^* end and BC from the 5*^′^* end with zero-mismatch tolerance (minimum overlap 18 bp); reads missing either primer were dropped. This removed a further 71 k reads, yielding 2.62 M primer-trimmed reads.

Syn1 transcripts are not natively polyadenylated, so any residual 3*^′^* A-run reflects the oligo-dT template-switch artifact and should be removed. Across the 2.62 M primer-trimmed reads the trailing-A run had median 30 nt and 99th percentile 34 nt, but a heavy tail reached 1.15 knt. Because the Syn1 genome contains no A-stretch longer than *∼*40 nt, runs *>* 70 nt cannot be biological and were discarded as likely concatemers or internal-priming artifacts; retained reads had up to 35 trailing As removed.

Cleaned reads were mapped to JCVI-Syn1.0 (CP002027.1) with minimap2 v2.30 [58] and sorted and indexed with samtools v1.23 [57], producing 2.62 M primary alignments. Alignments were generated with the -ax map-hifi preset, which is optimized for high-accuracy PacBio HiFi reads and enforces contiguous read-to-genome alignment without splice junction inference.

A per-read QC table was then built from the sorted BAM with pysam v0.22.1, grouping each read’s primary and supplementary alignments. Reads were failed if any of the following was violated: MAPQ *≥* 20; primary aligned fraction (Σ of M/I/=/X over query length) *≥* 0.70; primary soft-clip fraction *≤* 0.30; *|*query length *−* primary reference span*| ≤* 100 bp. Reads were additionally flagged as strong concatemers when they produced *≥* 2 supplementary alignments that either covered the same locus with a *≥* 1.5*×* reference-span ratio, or mapped to disjoint loci (different contig/strand, or *>* 200 bp genomic gap). All failing reads were removed from the final high quality (HQ) BAM. 99.3% of primary-mapped reads (2.62 M *→* 2.6 M) passed all QC thresholds and constitute the HQ

BAM used for all downstream analyses. Read length is typical of bacterial mRNA isoforms (median 1.56 kb; 5th–95th percentile 0.88–3.06 kb; 99th percentile 3.92 kb; maximum 10.9 kb). Mapping quality is uniformly high (median MAPQ 60, mean 60.0, minimum 20 after filtering). The primary aligned fraction is essentially saturated (median 1.00, mean 0.999, 5th percentile 0.996), and the soft-clip fraction is negligible (median 0, mean 0.12%, 95th percentile 0.41%). 95% of retained reads have *|*query length *−* reference span*| ≤* 10 bp and 99% *≤* 55 bp, confirming near-gapless full-length alignments. In the PacBio Iso-seq library, 0.03% of aligned bases fell within the two rRNA operons across all mapped reads, indicating effective rRNA depletion.

#### RNA Isoform Clustering

Isoform clustering of PacBio FLNC reads. 2.6 M primary FLNC alignments yielded 621 k unique (chrom, strand, pos5p0, pos3p0) tuples at single-bp resolution, with 77.3% supported by a single read — far exceeding the 911 annotated Syn1 genes and reflecting a combination of biological TSS/TTS micro-heterogeneity, alignment noise, and low-count degradation products.

A bare read-count threshold is unacceptably lossy: requiring n *≥* 10 reads per single-bp tuple discards *≈*874 k reads (*≈*34% of all primary reads), most of which sit on tuples directly adjacent to high-count tuples and represent biological wobble around a single underlying TSS/TTS rather than independent transcripts. Clustering is required to recover this signal for quantification and to deduplicate near-identical end positions for downstream analyses.

To choose a bp-tolerance (eps), cluster counts were swept over a range of tolerances on informative subsets (tuples with n *≥* 10 and n *≥* 50 reads). The cluster count decreased smoothly with increasing tolerance and showed an inflection between 10 and 15 bp before flattening beyond 20 bp. An eps of 10 bp was chosen, consistent with the typical end-position accuracy of PacBio CCS reads and the 10-bp deduplication tolerance used in the operon segmentation, and yielding biologically sensible counts (*≈*1.5 k distinct transcription units among *≈*900 genes organized in operons).

Tuples were clustered per (chromosome, strand) by complete-linkage under Chebyshev distance with a cutoff of eps = 10 bp. Complete linkage (rather than single linkage) was required to prevent chain-like merging through the dense carpet of low-count degradation-product tuples in highly expressed operons, which would otherwise fuse multiple real TSSs into a single giant cluster. No minimum-read filter was applied at cluster construction, so the isoform catalog is complete and downstream analyses select their own thresholds. For each cluster, the read-count-weighted median 5’/3’ end positions, the median absolute deviation (MAD) of member positions on each axis, and the total number of supporting reads were reported.

Complete-linkage clustering collapsed 621 k single-bp tuples into 267 k isoform clusters. The diameter guarantee was reflected in the empirical distributions: median MAD was 0 bp on both 5’ and 3’ ends, the 99th percentile was 5.0 bp (5’) and 4.5 bp (3’), and the maximum was bounded at 5 bp (= eps/2). Applying minimum-read thresholds to clusters rather than raw tuples substantially increased the quantified signal: at n *≥* 10, clustering retained *≈*21 k isoforms covering *≈*2.1 M reads (vs. *≈*17 k tuples covering *≈*1.7 M reads, +23%); at n *≥* 50, clustering retained *≈*4 k isoforms covering *≈*1.8 M reads (vs. *≈*3.3 k tuples covering *≈*1.5 M reads, +22%). The n *≥* 50 set was used for operon segmentation and the n *≥* 10 set for transcript visualization in subsequent figures.

Each cluster was annotated against the Syn1 gene model (GFF3) as sense, antisense, intergenic, or mixed using 95%-of-length assignment rules with a 50-bp minimum gene-overlap threshold.

#### Quantification of TPM from Sequencing Depth

Per-gene sense and antisense TPM were computed from the PacBio HQ per-strand depth tracks using the same depth-based definition applied to the Illumina data, namely the gene-length-normalized mean per-base depth scaled to transcripts per million against a single sense-plus-antisense denominator. Because the three technical replicates were pooled into a single library before mapping, no replicate averaging was applied, yielding one Syn1.0 PacBio TPM profile (available in Supplementary Data S2).

### Illumina Short-read Sequencing of Syn3A Transcriptome

#### Mapping Illumina Reads to Syn3A Genome

Raw Illumina reads (paired-end, 51 nt, *∼*12.7 M pairs per replicate) of Syn3A wild type at exponential phase from [10] were retrieved from NCBI SRA database. Per-base and per-read quality scores passed all checks and no Illumina adapters were detected, so no trimming was applied.

Reads were aligned to the JCVI-syn3A reference (NCBI accession CP016816.2) with bowtie2 v2.5.5 [56] in default paired-end mode. The overall alignment rate was 98.88%: 83.55% of pairs mapped concordantly exactly once, 10.43% concordantly with *>*1 hit (corresponding to the two ribosomal RNA operons of syn3A), and 6.02% of pairs were discordant or unpaired. SAM output was converted, sorted, and indexed with samtools v1.23 [57]. Strand-specific BAMs were generated using the dUTP / fr-firststrand flag convention (read 2 forward on + strand and read 1 reverse on + strand define plus-strand reads; their reverse complements define minus-strand reads), matching the Kapa Stranded RNA-Seq Kit library preparation, and per-strand sequencing-depth bedGraph tracks were computed with samtools depth.

#### Quantification of TPM from Sequencing Depth

Per-gene sense and antisense TPM were computed for Syn3A from the strand-specific depth tracks, using the same depth-based definition applied to the Syn1.0 Illumina data, namely a gene-length-normalized mean per-base depth scaled to transcripts per million against a single sense-plus-antisense denominator per dataset.

### Oxford Nanopore Direct-RNA Sequencing of Syn1 and Syn3A Transcriptomes

#### ONT RNA Sample Preparation

One mL aliquots of actively growing JCVI-syn1.0 and JCVI-syn3A cells were placed in 25 mL of SP4 + 17% KnockOut serum replacement in 50 mL conical tubes and incubated overnight at 37°C. The cells were harvested in mid- to late-exponential phase as indicated by a red-orange color of the phenol red pH indicator in the growth media, when the media pH was 7.0. Syn1.0 cultures achieved this growth stage in 16 hours and Syn3A cultures in 22 hours. Growth was arrested in the bulk cultures by placing the tubes on wet ice for 10 minutes. Keeping all tubes on ice, 1.4 mL of bulk culture was distributed into fifteen 1.8 mL microfuge tubes per culture and centrifuged at 4°C for five minutes at 10,000*×*g using a tabletop refrigerated centrifuge to pellet the cells. The supernatants from ten of the fifteen tubes were aspirated, and in the remaining five tubes the supernatant was used to suspend the pellets. The cell suspensions from those five tubes were then used to resuspend the pellets in five other tubes, and those more concentrated cell suspensions were used to resuspend the cells in the last five tubes, leaving both the Syn1.0 and Syn3A cells in five tubes of concentrated cell suspension. These tubes were centrifuged for five minutes at 4°C under 10,000*×*g, the supernatants were aspirated completely, and the remaining pellets were snap frozen using liquid nitrogen, transferred to dry ice, and immediately stored at -80°C.

Cellular RNA was extracted from Syn1.0 and Syn3A cell samples using a phenol-chloroform extraction protocol adapted from [55] and the ThermoFisher TRIzol^TM^ Reagent user guide. Briefly, 1 mL of 4°C TRIzol^TM^ was added to each tube containing frozen cell pellets, and the liquid was pipetted up and down multiple times to break up the cell pellet. The tubes were incubated at room temperature for 5 minutes to allow complete dissociation of nucleoprotein complexes. 200 *µ*L of chloroform was then added and the tubes were mixed by shaking for 30 seconds. After the tubes incubated at room temperature for 3 minutes, they were centrifuged at 12,000*×*g for 15 minutes at 4°C. The upper RNA-containing aqueous phase was pipetted without collecting any of the white interphase separating the aqueous from the phenol-chloroform lower phase, and placed in fresh 1.8 mL tubes already containing 500 *µ*L isopropanol. The tubes were incubated on ice for 10 minutes and then centrifuged at 12,000*×*g for 10 minutes at 4°C. The supernatant was aspirated and discarded off the white gel-like pellets, the tubes were inverted, and the pellets were allowed to air dry for 15 minutes. The RNA pellets were then dissolved in 20 *µ*L of RNase-free water containing 0.1 mM EDTA. RNA concentration was assessed using a Qubit RNA HS fluorometer assay and either an Agilent Bioanalyzer 2100 (Syn1.0) or an Agilent TapeStation 2200 (Syn3A). The next step was ribosomal RNA (rRNA) depletion. The original Syn1.0 sample was rRNA depleted using an Invitrogen RiboMinus^TM^ Bacteria 2.0 Tran-scriptome Isolation Kit. Briefly, 2 *µ*g of total Syn1.0 RNA was mixed with 100 *µ*L of RiboMinus^TM^ 1X hybridization buffer and 6 *µ*L of RiboMinus^TM^ Pan-Prokaryote Probe Mix, and the solution was brought up to 200 *µ*L using RNase-free water. The solution was denatured for 10 minutes at 70°C and then hybridized for 20 minutes at 37°C. The hybridization solution was mixed with RiboMinus^TM^ Magnetic Beads (1 mL of suspension) that had been washed in RNase-free water and resuspended in 400 *µ*L of RiboMinus^TM^ 1X hybridization buffer, and incubated at 37°C for 15 minutes. The sample tube was placed on a RiboMinus^TM^ magnetic stand, and the supernatant (600 *µ*L) containing the rRNA-depleted RNA was transferred to two new 1.8 mL tubes. To those tubes, each containing 300 *µ*L of RNA, premixed Binding Solution Concentrate (400 *µ*L) and Nucleic Acid Binding Beads (10 *µ*L) supplied with the kit were added and gently mixed. One mL of 100% ethanol was added to each tube, mixed, and incubated at room temperature for 5 minutes. The tubes were returned to the magnetic stand for 5 minutes, and the supernatant was aspirated and discarded without disturbing the beads. The beads were washed in 300 *µ*L of the kit Wash Solution, the tubes were returned to the magnetic stand, and the cleared supernatant was aspirated and discarded without disturbing the bead pellet. After the beads had dried for 5 minutes, 25 *µ*L of pre-heated (70°C) nuclease-free water was added and the tubes were incubated for 1 minute at room temperature to elute the RNA. The tubes were returned to the magnetic stand and the supernatant was carefully collected into a new microcentrifuge tube. The second Syn1.0 sample and the Syn3A sample were rRNA depleted using the NEBNext® rRNA Depletion Kit for bacteria. One *µ*g of RNA was brought up to 11 *µ*L with RNase-free water, and 2 *µ*L of NEBNext® Bacterial rRNA Depletion Solution and 2 *µ*L of NEBNext® Probe Hybridization Buffer were added. The reaction was heated to 95°C for 2 minutes and cooled to 22°C at a ramp rate of 0.1°C/sec. This was followed immediately by RNase H digestion, where 2 *µ*L of RNase H Reaction Buffer and 2 *µ*L of NEBNext Thermostable RNase H were added to the hybridized RNA solution to a total volume of 20 *µ*L, and incubated at 50°C for 30 minutes. Following a bead cleanup with Agencourt RNAClean XP beads (Beckman Coulter™, cat # A63987), the 15 *µ*L of the rRNA-depleted solution was poly(A) tailed at the 3*^′^* end.

The same poly(A) tailing protocol was used for Syn1.0 and Syn3A. RNA samples were mixed with a premade solution containing 2 *µ*L of 10X *E. coli* Poly(A) Polymerase Reaction Buffer, 2 *µ*L of ATP, and 1 *µ*L of *E. coli* Poly(A) Polymerase (New England Biolabs) and incubated at 37°C for 30 minutes. The reaction was stopped by adding EDTA to a final concentration of 10 mM. Following another bead cleanup along with the necessary quality control analysis, RNA was normalized to 9 *µ*L containing 140 ng for Syn1.0 and 50 ng for Syn3A RNA samples.

#### ONT RNA Sample Quality Control Analysis

An Agilent Bioanalyzer 2100 assay was used for assessing RNA quality and concentration of the first Syn1.0 sample both before and after rRNA depletion. For the second Syn1.0 and the Syn3A samples, both a Qubit RNA HS fluorometer assay and an Agilent TapeStation 2200 were used for assessing RNA concentration post-extraction from minimal cells. Initial Qubit values post RNA extraction were 3000 ng/*µ*L for the second Syn1.0 and 1890 ng/*µ*L for Syn3A. The Agilent TapeStation 2200 was used throughout the extraction and rRNA depletion steps to assess the quality of the RNA and depletion, giving an estimate of depletion through the absence of the 16S and 23S peaks (Supplementary Data S4). A Qubit RNA HS fluorometer was used to detect the final quantitation of RNA. Post poly-adenylation of RNA transcripts, the extent of DNA and buffer contamination in each of the RNA samples was tested by performing standard spectroscopic measurements (NanoDrop One) and using the Qubit 1*×* dsDNA HS assay kit (ThermoFisher Scientific). Input RNA samples for Oxford Nanopore Direct RNA Sequencing library preparation were finally quantified using the Qubit RNA HS assay kit, and RNA concentration values were used to normalize samples to the desired input volume of 9 *µ*L containing 50–500 ng of input RNA for library preparation.

#### ONT Library Preparation and Direct Sequencing

After normalization, libraries for Oxford Nanopore sequencing were prepared using the ONT Direct RNA Sequencing Kit (SQK-RNA002) with slight modifications. Specifically, 0.5 *µ*L of the RT adapter enzyme (reduced from 1 *µ*L) and 4 *µ*L of the RMX RNA adapter (reduced from 6 *µ*L) were used during library preparation for Syn1.0 and Syn3A. All samples were sequenced using R9.4 flow cells on the ONT GridION platform, with live base-calling enabled through the recommended MinKNOW (v23.07.5) scripts to generate FAST5 files. For Syn1.0, 819,435 reads were generated over two runs, and for Syn3A 734 k reads over a single run. In all run reports, base calls had 99.99–100% confidence.

#### Base-calling of Raw ONT Reads

Nanopore reads were collected as individual FAST5 files, which were combined into multi-read files for analysis, and current signals were converted to canonical bases by base-calling. For the first Syn1.0 sequencing, base-calling was performed with Guppy (v6.1.3; Oxford Nanopore Technologies, Ltd., 2022) using the RNA trim strategy and with calibration detection and filter enabled, on NVIDIA A5000 GPUs. For the second Syn1.0 replicate and the Syn3A experiments, Guppy (v7.0.9, GUI-based for the MinKNOW system) was used for base-calling with equivalent parameters.

#### Mapping ONT Reads to the Reference Genomes

Reads were retrieved from the NCBI SRA with the SRA Toolkit v3.4.1: Syn1.0 run 1 (SRR36199726, 234,477 reads), Syn1.0 run 2 (SRR36199725, 584,958 reads), and Syn3A (SRR36199724, 734 k reads). Direct-RNA reads deposited in the native 3*^′^ →*5*^′^* pore order (the Syn1.0 run 2 and the Syn3A run) were restored to 5*^′^ →*3*^′^* sense orientation by reversing the base order (seqkit v2.1.0, reverse only and not reverse-complement, since direct-RNA reads are single-stranded), whereas the Syn1.0 run 1 reads were already in sense orientation. Orientation was verified empirically: in the corrected orientation reads mapped sense to the annotated genes (over 97% of gene-overlapping reads), whereas the opposite orientation mapped almost nothing. Reads were aligned to the respective reference genome (JCVI-syn1.0, NCBI accession CP002027.1; JCVI-syn3A, CP016816.2) with minimap2 v2.30 [58] using the non-spliced long-read preset -ax map-ont -p 0.99 --MD, and sorted and indexed with samtools v1.23 [57]. The map-ont preset was used in place of the splice-aware preset because *M. mycoides* and its reduced derivative are intron-less, so N-skip (spliced) alignments would bridge tens to hundreds of kilobases of unrelated reference. Strandedness of the direct-RNA reads is preserved through the BAM flag bits, and per-strand sequencing-depth bedGraph tracks were computed with samtools depth, with forward-mapped reads defining the plus-strand transcripts.

For Syn1.0, run 1 yielded 207,480 primary mapped reads (88.5%; mean length 661 nt, mean Phred quality Q17.5) and run 2 yielded 456,755 primary mapped reads (78.1%; mean length 370 nt, Q31.7, 98.4% above Q20), the higher quality of run 2 reflecting its newer base-caller. For Syn3A, 559 k of 734 k reads (76.2%) mapped as primary alignments (mean length 383 nt, range 49–2,858 nt, mean quality Q31.2, 98.2% above Q20); the 175 k unmapped reads (23.9%) were of substantially lower quality (mean Q20.7, 61.0% shorter than 300 nt), consistent with the expected tail of truncated ONT direct-RNA molecules rather than reference mismatch. Up to 1.4% of primarily aligned bases fell within the two rRNA operons across the three Syn1.0 and Syn3A ONT datasets, indicating effective rRNA depletion.

#### Between-run Reproducibility and TPM Quantification

Per-gene sense transcripts-per-million (TPM) were computed from the strand-specific depth tracks using the depth-based definition applied to the Syn1.0 Illumina and PacBio data. The two Syn1.0 runs agreed only moderately (Pearson *r* = 0.82 on log_10_ TPM, with 36% of genes differing more than two-fold), the largest discrepancies being residual rRNA transcripts, which the RiboMinus^TM^-depleted run 1 retained far more abundantly than the NEBNext®-depleted run 2. Because the two runs used different depletion chemistries and differ substantially, they were treated as independent runs rather than averaged into a single quantitative profile, and the pooled alignment was retained only as a genome-browser coverage track. Each Syn1.0 run reproduced the Syn1.0 Illumina TPM profile with comparable fidelity (Pearson *r* = 0.61 and 0.62 on log_10_ TPM for runs 1 and 2), confirming that the direct-RNA data recover the same transcriptome as the short-read and PacBio platforms. The Syn3A ONT TPM profile was quantified in the same way and is reported alongside the Syn3A Illumina track (see the Syn3A short-read sequencing methods).

### Operon Identification from PacBio Transcriptomics of Syn1

#### Isoform-based operon segmentation

Of the 267 k clustered RNA isoforms, 4 k supported by *≥*50 reads were retained to suppress noise from sequencing artifacts and RNase degradation fragments. These isoforms were grouped into candidate operons by containment clustering: two isoforms were placed in the same cluster if one was fully contained within the other (within a 10 bp boundary tolerance), implemented via a union-find algorithm with a coordinate sweep line. This containment clustering collapses nested isoforms — the internal termination and RNA processing products of one operon — into the enclosing unit. This produced 313 initial operons (147 on the plus strand, 166 on the minus strand; median length 2,126 bp), which were then annotated against the 911 canonical *Syn1* genes, of which 304 operons contained at least one sense gene (104 single-gene, 98 two-gene, and 102 with three or more genes).

Adjacent same-strand operons can represent split fragments of a single operon, arising in two ways: either two operons that overlap (29 pairs, 24 sharing a gene), produced when RNase cleavage leaves a separate 5’ and 3’ piece, or two operons separated by a short gap (*≤*600 bp) that still contains a same-strand gene, a clustering coverage hole where no isoform passed the read threshold. Rather than resolving these by boundary location, each such candidate junction was tested for direct co-transcription from the read data: a pair was merged only when at least 50 strand-specific PacBio reads spanned the junction window (*±*80 bp) and the mean read depth across the junction was at least half that of the flanking regions, and was otherwise kept separate as a genuine terminator/promoter boundary. Of the 61 candidate junctions (29 overlapping, 32 gene-in-gap), 43 passed and 18 were kept separate; merges were applied pairwise with no transitive chaining (each operon joining at most one merge), producing 37 merged operons. A final union-find pass then merged any same-strand operons sharing identical sense gene loci, combining one further pair and yielding 275 isoform-derived operons.

After the initial segmentation, 215 of 911 genes (23.6%) remained uncovered by any isoform-derived operon. These were rescued through three complementary strategies: (i) 20 groups of consecutive uncovered genes (gap *≤*500 bp) were recovered by identifying spanning isoforms in the BAM file, (ii) the two rRNA operons (16S–23S–5S) were added manually, as their extreme secondary structure precludes faithful long-read capture, and (iii) the remaining 162 uncovered genes with non-zero spanning reads were added as single-gene transcription units. The final operon map comprised 459 operons (237 from isoform clustering, 37 merged, 1 gene-combined, 22 multi-gene rescue, and 162 single-gene rescue) and achieved complete sense-strand coverage of all 911 annotated *Syn1* genes (91.3% sense-only, 5.6% both sense and antisense, 3.1% antisense-only, 0% uncovered).

#### Locate Transcription Promoter and Terminator Signatures

Promoter and terminator signatures were characterized only for canonical operons, tentatively defined as isoform-derived operons (segmentation type isoform operon) whose transcription start site (TSS) and transcription termination site (TTS) both fall in intergenic regions, meaning that neither boundary lies inside a same-strand gene body. This restriction retains only operons whose mapped 5*^′^* and 3*^′^* ends reflect genuine promoter and terminator positions rather than RNase-truncated or internally nested transcript ends. The TSS was taken as the operon start0 on the plus strand and the end0 on the minus strand, with the TTS at the opposite boundary.

For each canonical TSS, the upstream promoter was read off the transcribed strand of the Syn1.0 genome (with circular wraparound). The *−*10 box was located by IUPAC consensus matching, scanning a 6-mer consensus TANAAT over the window from *−*12 to *−*7 and an extended 9-mer consensus TNNTANAAT over *−*15 to *−*7 relative to the TSS, each allowed a register shift of up to *±*2 bp, and selecting the offset that minimized the number of IUPAC mismatches (ties broken by the smallest shift). The *−*35 region (from *−*37 to *−*31) carried no fixed consensus and was instead summarized by a position-specific sequence logo, which showed it to be AT-rich. Information-content sequence logos were also generated for the *−*10 box and the TSS context. Rho-independent (intrinsic) terminators were predicted across the Syn1.0 genome with TransTermHP v2.09 [23], each prediction consisting of a stem-loop hairpin followed by a 3*^′^* poly-uridine tail and carrying a confidence score. A predicted terminator was assigned to a canonical operon as terminating when it fell within 50 bp downstream of the TTS in the transcription direction, or as internal when it lay inside the operon body at least 50 bp from the TTS. The terminator hairpins were rendered as sequence logos.

### RNA Processing Analysis

#### Ribonucleases in Syn1 and Syn3A

The ribonuclease complement of Syn1.0 was read from the genome annotation and assigned activities from the characterised *Bacillus subtilis* orthologues. Both Syn1.0 and Syn3A encode the same compact set: the endoribonucleases RNase III, RNase Y and RNase M5, the 5*^′^ →*3*^′^* exoribonucleases RNase J1 and RNase J2, the 3*^′^ →*5*^′^* exoribonucleases RNase R and YhaM, and the *trans*-translation factors tmRNA and SmpB (Table S3) [26, 38]. Per-cell abundances were the proteomic copy numbers (Section on proteomics), and substrate signatures were taken from the corresponding *B. subtilis* enzymes.

#### RNA-processing endpoint analysis

Post-transcriptional processing was quantified from the clustered PacBio isoforms. Isoforms supported by at least ten reads were retained, giving 20,885 isoforms. A per-strand mask of protein-coding gene bodies was built from the Syn1.0 GFF3 annotation, and the 5*^′^* and the 3*^′^* genomic endpoints of each isoform were labelled intragenic when fell inside a gene body and intergenic otherwise. Each isoform was assigned to one of four end-context categories — unprocessed (both ends intergenic), 5*^′^*-eroded only, 3*^′^*-eroded only, or both-eroded — and the composition was tabulated both by isoform count and by read. The erosion asymmetry was summarised as the ratio of the read weight carried by 5*^′^*-intragenic to that carried by 3*^′^*-intragenic isoforms. Independently, every annotated start and stop codon fully contained within an isoform was counted, and a reading frame that contained its start codon but not its stop codon was scored as 3*^′^*-truncated. For the single-gene panels, the isoforms contained within an operon were coloured by end-context and ordered into the erosion ladder; for the ATP-synthase panel the member isoforms of the two segmented operons were coloured by transcription block.

#### Homology Comparison of RNase III and RNase Y between B. subtilis and Syn1

RNase III and RNase Y, the endoribonucleases that carry out most intra-operonic cleavage, were compared between *B. subtilis* (NCBI NC 000964.3) and Syn1.0 to confirm conserved catalytic and RNA-binding architecture. Domain organisation was assigned with InterPro, and pairwise coverage and identity were taken from the protein alignment. Both enzymes retained their full *B. subtilis* domain architecture in Syn1.0: RNase III (249 versus 232 residues) kept its RNase III and double-stranded-RNA-binding domains at 91% coverage and 45% identity, and RNase Y (520 versus 509 residues) kept its RNase Y, KH and HD domains at 98% coverage and 35% identity.

### Novel Transcription and Translation in Syn1

#### Anti-sense and Intergenic Transcription

Each PacBio isoform cluster was labelled base-by-base against the canonical Syn1.0 gene model, and its sense, antisense, and intergenic coverage fractions were computed. Antisense coverage was carried by 1.4% of the isoforms (267 of those with *≥* 10 supporting reads), which collapsed to 89 representative clusters after merging near-identical ends within 10 bp. These were classified by the arrangement of their sense and antisense segments into three cases: antisense-initiated transcripts from a spurious promoter in the AT-rich genome (anti *→ . . .*), sense transcripts reading through into an antisense gene at an operon boundary (sense *→* anti), and read-through transcripts with an antisense gene embedded between sense genes (sense *→* anti *→* sense). Spurious-promoter transcripts accounted for 59 of the 89 clusters (66%), and read-through transcripts, including 4 embedded cases, for the remaining 30 (34%).

#### Novel Open-Reading Frames from Full-length RNA Isoforms

The abnormal isoforms, defined as those with *>*10 supporting reads and an abnormal fraction (*f*_antisense_ + *f*_intergenic_) *>* 0.2, gave 837 isoforms as input to the ORF-prediction pipeline.

All candidate start codons within the 837 isoforms were evaluated using OSTIR v1.1.2 [31], which predicts translation initiation rates (TIR) by modelling Shine–Dalgarno (SD) complementarity to the Syn1.0 16S rRNA anti-SD (ACCUCCUUU) and local mRNA secondary structure near each AUG. For every predicted start site, the downstream reading frame was extended to the first in-frame stop codon (TAA or TAG) under genetic code 4, producing *∼*29,000 candidate ORFs (*≥*15 amino acids).

ORFs overlapping known sense reading frames were excluded, and the remainder were ranked by a synthesis rate score (isoform read count *×* TIR). The top 100 ORFs (of which 38% exhibited strong SD-mediated initiation) were deduplicated by amino acid sequence to 48 unique peptides and subjected to *in silico* trypsin digestion (one missed cleavage; peptides of 7–25 residues). Comparison of the resulting tryptic peptides against the canonical Syn1.0 proteome identified 47 novel ORFs producing at least one proteotypic peptide suitable for detection by bottom-up mass spectrometry. These novel ORFs were appended to the Syn1.0 protein database, and the proteomics raw data were re-searched with Spectronaut.

### Proteomics of Syn1 and Syn3A

#### Sample Preparation – Cell lysis and Protein digestion

Cell pellets of Syn1.0 and Syn3A strains were mixed with 100 *µ*L of SDS-based lysis buffer (4% SDS, 100 mM Tris-HCl, pH 8.0) containing final concentration of 10 mM Tris (2-carboxyethyl) phosphine (TCEP) and 40 mM chloroacetamide (CAA), vortexed for 5 min, and then boiled at 95°C for 10 min. The lysate samples were then processed for proteomics analysis following the E3technology procedure reported recently [59]. In brief, the lysates were mixed with 4*×* volume of 80% acetonitrile (ACN) and then transferred to E3filters (CDS Analytical, Oxford, PA) followed by centrifugation at 3,000 rpm for 2 min. The filters were washed with 500 *µ*L of 80% ACN for three times, and then transferred to clean collection tubes. For protein digestion, around 1.0 *µ*g of trypsin and 200 *µ*L of digestion buffer (50 mM triethylammonium bicarbonate, TEAB) were added to each sample followed by incubation at 37°C for 16-18 hours with gentle shaking (300 rpm). After digestion, the filters were first spun at 3,000 rpm for 2 min to collect the flow through, and then eluted sequentially with 200 *µ*L of 0.2% formic acid (FA) in water and 200 *µ*L of 0.2% FA in 50% ACN. The pooled elution was dried in SpeedVac, desalted with C18 StageTips, and stored under -80°C until further analysis.

#### LC-MS/MS analysis and protein identification

The LC-MS/MS analysis was performed using an UltiMate 3000 RSLCnano system in conjunction with an Orbitrap Eclipse mass spectrometer and FAIMS Pro Interface (Thermo Scientific), similar to the procedures described recently [60]. In brief, the peptides resuspended in LC buffer A (0.1% FA in water) were first loaded onto a trap column (PepMap100 C18,300 *µ*m *×* 2 mm, 5 *µ*m particle, Thermo Scientific), and then separated on an analytical column (PepMap100 C18, 50 cm *×* 75 *µ*m i.d., 3 *µ*m; Thermo Scientific) at a flow of 250 nL/min. A linear LC gradient was applied from 1% to 25% mobile phase B (0.1% formic acid in acetonitrile) over 125 min, followed by an increase to 32% mobile phase B over 10 min. The column was washed with 80% mobile phase B for 5 min, followed by equilibration with mobile phase A for 15 min. For MS analysis, the Detector type was Orbitrap with resolution of 120,000; Precursor MS range (m/z) was 380–980; AGC target was Standard; Maximum injection time mode was Auto. For data-independent acquisition (DIA) MS/MS analysis, the Isolation mode was Quadrupole; DIA Window type was Auto and Isolation Window (m/z) was 8 with an overlap of 1; activation type was HCD with fixed collision energy mode (30%); the Detector Type was Orbitrap with a resolution of 30,000; Normalized AGC target (%) was 800, and the Maximum injection time mode was Auto; the Loop Control was 2 sec. For FAIMS compensation voltages (CV) setting, a 3-CV combination (-40, -55, and -75) was applied.

For proteome identification, the MS raw data were processed using Spectronaut software (version 19), a library-free DIA analysis workflow with directDIA+ and the in-house curated protein databases (Syn1.0, 828 sequences; Syn3A, 458 sequences). Detailed parameters for Pulsar and library generation include: Trypsin/P as specific enzyme with minimum peptide length of 7 amino acids and maximally 2 missed cleavages; toggle N-terminal M turned on; Oxidation on M and Acetyl at protein N-terminus as variable modifications; Carbamidomethyl on C as fixed modification; FDRs at PSM, peptide and protein level all set to 0.01; Quantity MS level set to MS2, and cross-run normalization turned on. The MS raw data associated with this study has been deposited onto the MassIVE Repository with the accession MSV000099558.

#### Relative Protein Quantification

Protein abundances were quantified from the Spectronaut iBAQ (intensity-based absolute quantification) values, which serve as a within-sample proxy for molar protein amount. For each of the three replicates per organism, iBAQ was converted to a relative abundance in parts per million (iPM), iPM*_i_* = 10^6^ iBAQ*_i_/* Σ*_j_* iBAQ*_j_* , and the per-protein mean iPM across replicates was used for all downstream analyses, with the across-replicate coefficient of variation retained as a quality metric. For Syn1.0, 721 of the 828 annotated protein-coding genes (87.1%) carried a quantified iPM. For Syn3A, 446 of the 455 annotated protein-coding genes (98.0%) carried a quantified iPM. The per-gene relative abundances (iPM) are available in Supplementary Data S2.

#### Absolute Protein Quantification

The Syn1.0 mass spectrometry data provides relative abundances for 721 of the 828 protein-coding genes, which can be converted to absolute abundances by scaling the relative amount with the total number of proteins in the cell. The total number of proteins has not been measured directly within Syn1.0, but can be estimated via an approach similar to that used in [2]. First the dry mass of the cell and protein mass fraction are needed (all parameters used are listed in Table S4). The protein mass fraction is assumed to be 58.2%, which was measured in the parent organism, *M. mycoides* [34]. The dry mass of the cell, 12.8 fg, was estimated based on parent organism *M. mycoides*, which is similar to the dry mass of Syn3A as in our previous publication [2]. Accordingly, the average JCVI-syn1.0 cell has approximately 127,000 proteins; the cell volume is calculated as the sphere of 220 nm radius, giving 0.044 *µm*^3^ [11]. Thus, a protein density of 2.87 *×* 10^6^ *proteins/µm*^3^ is estimated, which is in rough agreement with 3.5 *−* 4.4 *×* 10^6^ *proteins/µm*^3^ measured in *E. coli* [61]. Finally, absolute protein abundances were calculated using the following formula:

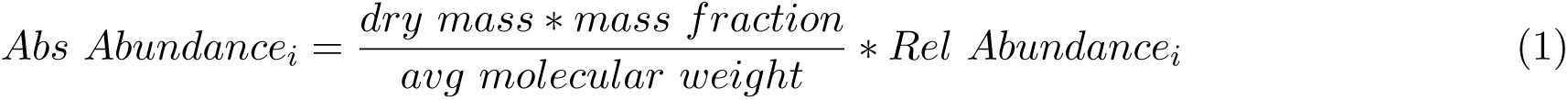

**Table S4:** Parameters for absolute protein copy-number quantification of Syn1.0 and Syn3A.

| Parameter | Value | Unit | Source |
| --- | --- | --- | --- |
| Avogadro constant ($N_A$) | $6.022 \times 10^{23}$ | $\text{mol}^{-1}$ | — |
| Syn1.0 protein mass fraction | 58.2 | % | [34]<sup>a</sup> |
| Syn1.0 cell diameter | 439 | nm | [11] |
| Syn1.0 cell volume | 0.044 | $\mu\text{m}^3$ | This study |
| Syn1.0 total cellular dry mass | 12.8 | fg | [34]<sup>b</sup> |
| Syn1.0 iPM-weighted mean protein MW | 35.2 | kDa | This study |
| Syn1.0 total proteins per cell | 127,800 | — | This study |
| Syn1.0 protein number density | $2.87 \times 10^6$ | $\mu\text{m}^{-3}$ | This study |
| Syn3A protein mass fraction | 54.727 | % | [2] |
| Syn3A cell diameter | 400 | nm | [2] |
| Syn3A cell volume | 0.034 | $\mu\text{m}^3$ | This study |
| Syn3A total cellular dry mass | 10.2 | fg | [2] |
| Syn3A iPM-weighted mean protein MW | 33.4 | kDa | This study |
| Syn3A total proteins per cell | 100,300 | — | This study |
| Syn3A protein number density | $3.0 \times 10^6$ | $\mu\text{m}^{-3}$ | This study |

Absolute copy number per protein = (dry mass × protein mass fraction) / mean molecular weight × relative abundance (iPM/ $10^6$ ), following [2]; see Equation 1. The iPM-weighted mean molecular weight and total proteins per cell are derived from the proteome of each organism.
<sup>a</sup>Measured in the parent organism *Mycoplasma mycoides* subsp. *capri*, which has similar genome size as Syn1.0.
<sup>b</sup>Estimated based on the parent organism *Mycoplasma mycoides* using the same method as in [2]. A density of 1.1 g/ml for the Syn1.0 and Syn3A cells was assumed [35]; and around 4.8 $\mu\text{l}$ water/mg cellular protein as measured in *Mycoplasma mycoides* capri serovar capri PG3 [36], corresponding to a cellular water content of 72%.

The resulting per-gene absolute copy numbers are available in Supplementary Data S2.

The same estimation was applied to Syn3A using its independently determined cellular parameters (Table S4). The Syn3A protein mass fraction, 54.727%, and total cellular dry mass, 10.2 fg, were taken directly from the whole-cell metabolic reconstruction of Syn3A [2], and the cell volume from a sphere of 200 nm radius (0.034 *µm*^3^). Of the 455 annotated Syn3A proteins, 446 carried a quantified iPM, and Equation 1, evaluated with an iPM-weighted mean molecular weight of 33.4 kDa, yielded approximately 100,000 proteins per average cell. The corresponding protein density, 3.0 *×* 10^6^ *proteins/µm*^3^, is close to the Syn1.0 estimate. Because Syn1.0 and Syn3A were quantified with independently determined dry masses and protein mass fractions, their absolute copy-number scales differ (Results). The resulting per-gene Syn3A absolute copy numbers accompany the Syn3A proteome annotation. The parameters used for both organisms are listed in Table S4 (placed with the correlation supplementary figure).

#### Localization of Proteome

Subcellular localization was predicted from sequence for every Syn1.0 protein, since membrane-associated proteins are systematically under-recovered by the trypsin-digestion workflow and are therefore handled separately in the transcriptome–proteome correlation. Transmembrane regions (TMRs) were predicted with DeepTMHMM [62] and signal peptides with SignalP 6.0 [63]. Each protein was assigned to one of four classes by priority: a SignalP secretory or lipoprotein signal peptide marked it as extracellular or lipoprotein, respectively; otherwise one or more DeepTMHMM TMRs marked it as membrane, and proteins with no TMR were assigned cytoplasmic. Of the 721 detected Syn1.0 proteins, 516 were cytoplasmic, 126 membrane, 68 lipoprotein, and 11 extracellular. Median copy number per cell fell steeply with membrane association, from 47 for cytoplasmic proteins to 21 (lipoprotein), 10 (membrane), and 3 (extracellular), consistent with the under-digestion and under-recovery of membrane-embedded proteins rather than a true abundance difference. The per-protein localization assignments are available in Supplementary Data S2.

### Correlation Between Transcriptome and Proteome of Syn1 and Syn3A

Gene-level proteome abundance (relative iPM) was correlated against transcriptome abundance (sense TPM) to test how faithfully transcript level predicts protein level in Syn1.0, and the deviations from this relationship were then attributed to translation effects. The two layers were joined by locus tag, and the correlation was restricted to the 717 mRNA genes with non-zero signal in both. Illumina sense TPM was used as the transcriptome axis, following the convention for short-read quantification. On a log_10_ scale, transcript and protein levels were positively correlated across all detected proteins (Pearson *r* = 0.61, *r*^2^ = 0.38, *n* = 717) and more tightly for the cytoplasmic subset (Pearson *r*= 0.70, *r*^2^ = 0.49, *n* = 512), consistent with the under-recovery of membrane, lipoprotein, and extracellular proteins by the digestion workflow. For each gene the residual was defined as the offset of log_10_(iPM) from the ordinary-least-squares fit on log_10_(TPM), so that a positive residual marks more protein than the transcript predicts (efficient translation or a long-lived protein) and a negative residual marks less. Translation initiation and translation elongation were evaluated for their ability to explain this residual. The combined per-gene transcriptome and proteome table, together with the predicted TIR and CAI, is available in Supplementary Data S2.

#### Translation Initiation Rate Prediction

Per-gene translation initiation efficiency was predicted from sequence to test whether initiation rate accounts for the transcriptome–proteome residual. For each detected mRNA gene, the PacBio isoforms (read count *≥* 10) whose transcribed span covered the full ORF were identified, and an initiation window spanning 30 nt of 5*^′^* UTR (or the available UTR, whichever was shorter) through 30 nt into the coding sequence was extracted from the Syn1.0 genome with circular wraparound. Each window was scored with OSTIR v1.1.2 [31] using the Syn1.0 16S rRNA anti-Shine–Dalgarno tail (5*^′^*-ACCUCCUUU-3*^′^*), taking the prediction at the annotated start codon, and the translation initiation rate (TIR) together with its thermodynamic sub-terms (Shine–Dalgarno, standby-site, spacing, and 5*^′^* mRNA-folding Δ*G*) was recorded per isoform.

A read-count-weighted mean TIR was then computed for each gene, yielding a valid TIR for 563 genes. The contribution of initiation was quantified by adding log_10_(TIR) to the baseline ordinary-least-squares model log_10_(iPM) *∼* log_10_(TPM) (baseline *r* = 0.60, *R*^2^ = 0.35 on this subset) and by correlating log_10_(TIR) against the baseline residual. Translation initiation explained little of the residual: it raised the model *R*^2^ by only 0.019 (1.9% of the variance), and log_10_(TIR) correlated weakly with the residual (Pearson *r* = 0.17, *P* = 5 *×* 10*^−^*^5^). Dissecting the OSTIR sub-terms against the residual, the Shine–Dalgarno hybridization Δ*G* carried the strongest association, though it remained weak (Pearson *r* = *−*0.12, *P* = 4.2 *×* 10*^−^*^3^; more negative Δ*G*, i.e. stronger pairing, associated with a slightly higher residual), while the standby-site (*r* = *−*0.02, *P* = 0.7), spacing (*r* = *−*0.07, *P* = 0.08), and 5*^′^* mRNA-folding (*r* = 0.08, *P* = 0.06) terms carried essentially none.

#### Codon Adaptation Index (CAI)

Codon optimality was quantified as a proxy for translation elongation efficiency and tested against the residual. A codon adaptation index (CAI) [33] was computed for all 828 frame-valid Syn1.0 coding sequences under the Mycoplasma genetic code (UGA = Trp), with the reference set of highly expressed genes taken as the top 20% of protein-coding genes by mean iPM (144 of the 721 detected genes, 91% of them cytoplasmic). CAI was essentially independent of transcript level (CAI vs log_10_(TPM), Pearson *r* = 0.006, *P* = 0.87) yet correlated with the proteome residual (Pearson *r* = 0.36, *P* = 4 *×* 10*^−^*^23^), carrying information nearly orthogonal to the transcriptome term. Adding CAI to the baseline model raised the explained variance from *R*^2^ = 0.38 to 0.46 (Δ*R*^2^ = +0.080, a 21% gain over the baseline; *n* = 717), and the improvement held within the cytoplasmic subset (*R*^2^ from 0.49 to 0.54, Δ*R*^2^ = +0.052; *n* = 512).

#### Repeating the analysis in Syn3A

The same transcriptome–proteome comparison was carried out for Syn3A, joining Illumina sense TPM to relative iPM by locus tag over the 446 mRNA genes detected in both layers. Protein localization, annotated for Syn3A on a finer scheme, was collapsed onto the four Syn1.0 classes: trans-membrane proteins were treated as membrane, peripheral-membrane and unassigned proteins were grouped with the cytoplasmic class, and the lipoprotein and extracellular classes were retained. Transcript and protein levels correlated to a similar degree as in Syn1.0 (Pearson *r* = 0.63 across all detected proteins, *n* = 446, and *r* = 0.65 for the cytoplasmic subset, *n* = 352).

Translation initiation was re-predicted from scratch, because Syn3A lacks a long-read isoform set. With no isoforms to define transcript ends, OSTIR was run at the gene level on a fixed window spanning 30 nt upstream through 30 nt into the coding sequence at each annotated start codon, using the same anti-Shine–Dalgarno tail (ACCUCCUUU) but without the read-count weighting applied in Syn1.0. As in Syn1.0, initiation explained little of the residual (Δ*R*^2^ = +0.015, with log_10_(TIR) versus the residual at Pearson *r* = 0.15).

The codon adaptation index was recomputed with Syn3A’s own reference set of highly expressed genes (the top 20% by iPM, 89 of 446 genes), since that reference set defines the codon weights. CAI again carried post-transcriptional information, correlating with the residual (Pearson *r* = 0.46) and raising the explained variance from *R*^2^ = 0.39 to 0.52 (Δ*R*^2^ = +0.127, *n* = 446), a somewhat larger gain than in Syn1.0.

### Genome Reduction

The transcriptional consequences of minimization from JCVI-syn1.0 to JCVI-syn3A were dissected with a fixed pipeline that first localized every deletion on a common coordinate frame, then propagated each deletion through Syn1.0’s operon structure, and finally tested the predicted regulatory effects against Syn3A reads. All coordinates were handled as 0-based half-open intervals on the circular chromosome, with neighbor and intergenic-distance calculations wrapping at the origin.

#### Whole-genome alignment of Syn3A onto Syn1.0 and deletion extraction

The Syn3A genome (CP016816.2, 543,379 bp) was aligned onto the Syn1.0 genome (CP002027.1, 1,078,809 bp) to recast minimization as a set of discrete, locatable deletions on a single reference frame. Alignment used nucmer from MUMmer v3.23 [64] in --maxmatch mode (minimum cluster length 100 bp, minimum exact seed 20 bp), and the delta was reduced to a single best one-to-one mapping with delta-filter (-1; *≥* 95% identity, *≥* 100 bp). Single-base differences and rearrangement breakpoints were tabulated with dnadiff and show-snps. Deletions were defined as Syn1.0 intervals covered by no one-to-one alignment block: the blocks were merged across gaps *≤* 50 bp and complemented against the Syn1.0 genome with BEDTools v2.30.0 [65], and complement intervals shorter than 50 bp were discarded as boundary noise. This yielded the authoritative set of 95 deletions (*≥* 50 bp) removing 536,543 bp (*∼*536 kb) of Syn1.0 sequence, each annotated with the Syn1.0 genes it overlapped by intersection against the Syn1.0 gene annotation. Across the retained backbone the two genomes were 99.90% identical, differing by only 36 single-nucleotide polymorphisms (SNPs) and 12 short insertions or deletions (indels), with no inversions and no translocations, and gene order was preserved apart from a single relocated block (*lap*/0154), so expression differences at retained genes are not attributable to sequence divergence. Locus tags correspond one-to-one between organisms with the numeric suffix preserved (MMSYN1 NNNN *↔* JCVISYN3A NNNN), which was used throughout to map Syn1.0 genes to their Syn3A orthologs.

#### Overlaying deletions onto Syn1.0 operons

Each of the 95 deletions was intersected at single-base resolution with the 459 Syn1.0 operons annotated from PacBio long reads (Supplementary Data S1). Every operon was described along two orthogonal axes. The truncation axis classified each overlapping deletion on its own as fully deleted (both operon bound-aries removed), 5’ truncation gene or 5’ truncation UTR, 3’ truncation gene or 3’ truncation UTR (a boundary cut that reached a sense-gene coding sequence, CDS, or stopped in the untranslated region, UTR, respectively), or intra truncated (lying strictly inside the operon, neither boundary touched). The 5*^′^* versus 3*^′^* assignment was strand-aware, and an operon hit by more than one deletion took the sorted union of its per-hit cases. By this span-level classification 181 operons were fully deleted and 162 untouched, with boundary truncations reaching a coding sequence more often at the 3*^′^* end (47 operons) than the 5*^′^* end (28), in agreement with the gene-level pattern. The gene-deletion axis classified the operon by which of its sense genes were lost as intact, all deleted, leading deleted, lagging deleted, or intra deleted. By this axis 235 operons lost all of their sense genes and 172 were intact, leaving 52 partially deleted (21 leading-edge, 20 lagging-edge, 11 internal). The resulting per-operon table (sheet Operon classification of Supplementary Data S3) was the structural input to the junction, co-transcription, and gene-impact analyses below.

#### Deletion-junction taxonomy

Every deletion was reframed as a junction between the two retained operon fragments it joins. Operon L was the Syn1.0 operon of the nearest Syn3A-retained gene to the left of the deletion and operon R the nearest retained gene to the right; genes absent from Syn3A were skipped, and a gene covered only in antisense by an operon was assigned to that operon and treated as transcribed on the operon’s strand. The relative orientation of the two flanking operons defined strand relationship: intra operon (operon L = operon R, the deletion internal to one operon), tandem (the two operons co-oriented, the only configuration in which one transcript can run across the junction), convergent (operon L on + and operon R on *−*, their terminators facing the gap), and divergent (operon L on *−* and operon R on +, their promoters facing the gap). At a tandem junction only two regulators decide the outcome, the upstream operon’s terminator (its 3*^′^* end facing the gap) and the downstream operon’s promoter (its 5*^′^* end facing the gap), each scored as lost when its coordinate fell inside the deletion. The four combinations defined junction type: clean excision (both regulators kept, whole operon(s) excised between independent neighbors), decapitation (downstream promoter lost, predicting an expression drop), readthrough extension (upstream terminator lost, predicting additive read-in), and fusion (both lost, predicting a single chimeric transcription unit). Of the 95 junctions, 53 were tandem, 19 convergent, 15 divergent, and 8 intra-operon; the 53 tandem junctions comprised 30 clean excisions, 11 read-through extensions, 9 decapitations, and 3 fusions. A consistency check against the operon-overlay classification was run, mapping each lost regulator to its expected truncation side (a lost promoter to a 5*^′^* truncation, a lost terminator to a 3*^′^* truncation) and flagging disagreements (sheet Deletions of Supplementary Data S3).

#### Co-transcription tests from Syn3A reads

Whether a same-strand gene pair was still co-transcribed in Syn3A was tested directly from reads, using the Syn3A ONT direct-RNA alignment and the Illumina per-strand sequencing depth (pysam v0.22.1, matching-strand depth throughout). A pair passed the strict, spanning test when at least one primary ONT read on the matching strand fully enclosed both CDSs and the intergenic region between them. A pair passed the loose, bridging test when the number of ONT reads that overlapped each CDS by *≥* 50 bp and covered the intergenic gap reached max(1, min(10, 0.20 *× d*_min_)), where *d*_min_ is the lower of the two mean ONT gene depths, and when the intergenic gap was either shorter than 10 bp or carried mean Illumina depth *≥* 20% of the lower flanking-gene Illumina depth. Operon-internal co-transcription walked every consecutive retained sense-gene pair within an operon and called the operon preserved only when all of its pairs passed; the operons tested were routed from the junction table into pristine controls (gene-deletion pattern intact and flanking no junction, so the reduction never touched them) and intra operon operons (a deletion removed interior genes). Cross-junction co-transcription tested the newly adjacent gene pair at each inter-operon junction, each gene mapped to its Syn3A ortholog by locus suffix, stratified by junction type: fusion junctions were predicted to co-transcribe and clean excision junctions served as the negative control, while convergent and divergent junctions lie on opposite strands and were reported as structural negatives. By these tests fusion junctions co-transcribed in 2 of 3 cases against 3 of 30 clean-excision controls, on a pristine single-operon baseline of 27 of 45 operons. Because Syn3A ONT depth is low, most positive calls rested on the loose bridging criterion rather than on fully spanning reads (sheet Coexpression of Supplementary Data S3).

#### Per-gene transcriptional-impact classification

Each Syn1.0 gene retained in Syn3A (no base of its CDS overlapping any deletion) was annotated with its same-strand transcription-direction neighbors, its clockwise and counter-clockwise neighbors in both genomes, and the length of contiguous unaltered context flanking it (Syn1.0 frame, circular). Each retained gene was then assigned a gene impact class describing how the reduction changed the source of its transcription, integrating the operon-overlay, junction, and co-transcription results. Classes were assigned by precedence, the first matching rule winning: promoter lost (the operon lost its 5*^′^* promoter with no replacement, assigned at operon level to all of its retained genes), promoter disconnected (the promoter was kept but the operon split internally, separating downstream genes from it), new promoter fusion (the operon lost its own promoter but fused to the upstream operon’s promoter, confirmed by cross-junction co-transcription), readthrough exposed (the gene kept its promoter while the upstream operon lost its terminator), promoter proximity changed (an intra-operon deletion removed interior genes while the operon stayed co-transcribed), context only (the operon’s own 3*^′^* structure was truncated but its promoter and upstream context were intact), and unaffected (the operon’s own structure was intact, including operons that merely neighbor a deletion). Of the retained genes, 360 were unaffected, 45 context only, 42 promoter lost, 24 readthrough exposed, 17 promoter proximity changed, 6 promoter disconnected, and 3 new promoter fusion. This classification is purely structural; the corresponding changes in transcript and protein abundance were quantified separately (sheet Gene table of Supplementary Data S3, with the abundance comparison described in the following subsection).

#### Relative-abundance normalization and the paired expression-change table

Transcript and protein abundances were compared between the two organisms after a normalization that removes the artefact of the reduced gene complement. TPM and iPM are both relative units that each sum to 10^6^ within an organism, so when several hundred genes are absent from Syn3A the same fixed pool is divided among fewer genes and every retained gene’s value is mechanically inflated, leaving a naive Syn3A/Syn1.0 ratio well above 1 even for genes whose true abundance is unchanged. To remove this baseline, each gene’s value was expressed relative to the per-gene mean of its own organism and subset, rel*_i_* = *v_i_/v̄*, where *v̄* is the mean over the genes detected in that subset (coding or non-coding), so that the average gene equals 1 in both organisms and an unchanged gene gives a fold change of *∼*1 regardless of the gene-count difference. The same normalization was applied independently to the transcriptome and the proteome, yielding relTPM and relIPM, and two change metrics were derived for each layer: a fold change, the Syn3A relative value divided by the Syn1.0 relative value, and an absolute change, their difference, which is robust where a denominator is small. Illumina TPM was used for both organisms because ONT direct-RNA is unreliable for quantification, with ONT retained only for quality-control correlation against the Illumina tracks; the non-coding subset (rRNA and tRNA) was the sole exception and took Syn3A ONT TPM, since the Ribo-Zero Illumina library detects none of these species. The proteome layer used the mean-normalized iPM from the relative protein quantification described above, carrying the same fold-change and absolute-change definitions. The transcriptome and proteome layers were joined by locus number into a single paired table (compiled into sheet Gene table of Supplementary Data S3), from which the protein-specific figures and outlier reports were generated separately.

#### Deleted-gene occupancy and function-category reallocation

The expression capacity freed by minimization was quantified as the share of Syn1.0’s transcriptome and proteome carried by loci absent from Syn3A, using raw (un-normalized) Syn1.0 TPM and iPM so the values read as fractions of the whole, with the deleted loci partitioned by RNA type (sheet Summary stats of Supplementary Data S3). The reorganization of the retained pool was then analyzed by curated function, joining coding genes to their Primary, Secondary, and Tertiary function annotation by locus number. Cross-organism category shares require an additional correction, because Syn1.0’s full-pool mean is diluted by the deleted genes and therefore inflates the relative values of retained genes relative to Syn3A, which consists entirely of retained genes. The Syn1.0 side was therefore renormalized to the retained-gene pool before any category comparison, dividing each Syn1.0 relative value by the mean over detected genes that are kept in Syn3A, so that retained Syn1.0 genes again average 1 and are matched to Syn3A; the genome-wide fold-change distributions and the per-gene impact-class comparison retained the original full-pool normalization, whereas every per-category share used this deletion-corrected basis. For each category the mRNA-pool share in each organism and its change, the median fold and absolute change, and a two-sided Mann–Whitney *U* test of log_10_ fold change against all remaining genes were reported for categories with at least three detected genes (sheet Function categories of Supplementary Data S3).

#### Macromolecular-complex abundance

The assembled abundance of two complexes whose subunits are all protein-coding, RNA polymerase (RNAP) and the degradosome, was estimated from a limiting-subunit rule whereby surplus subunits are turned over, so that the complex copy number tracks its lowest stoichiometry-adjusted component (the minimum over subunits). Core RNAP was estimated as the minimum of *rpoA*/2, *rpoC*, and *rpoB* (the *α*_2_*ββ^′^* core, with *σ* factors excluded), and the degradosome as the minimum of *rny*, *rnjA*, and *yhaM* + *rnr* [29]. Each estimate was computed in both relTPM and relIPM for each organism, the Syn3A/Syn1.0 ratio being the interpretable cross-organism quantity, and related transcription and RNA-turnover capacity to Syn3A’s longer cell cycle (sheet Summary stats of Supplementary Data S3).

## Acknowledgements

E.F., T.A.B., Z.R.T., J.I.G. and Z.L.-S. were supported by NSF MCB 2221237. A.P.M., J.E.C. and Y.-l.G. were supported by the National Institute of General Medical Sciences of the National Institutes of Health under Award Number R01GM139949. E.F., A.T.A., A.P.M., J.I.G. and Z.L.-S. were also supported in part by the NSF Science and Technology Center for Quantitative Cell Biology (NSF DBI 2243257).

The authors thank Pratap Venepally (JCVI Rockville) for informative discussion about the ONT experiments and data analysis. The authors thank Alvaro Hernandez (Roy J Carver Biotechnology Center at UIUC) for carrying out the Illumina and PacBio sequencing of Syn1.0. The authors thank Rasmus Jensen (The European Molecular Biology Laboratory) for the preliminary analysis of tomograms of Syn3A. We also thank Kim Wise (JCVI La Jolla), James Pelletier (Centro Nacional de Biotecnología), and Benjamin Gilbert (UIUC) for informative discussions.

## Data availability

The raw sequencing data generated in this study are deposited in the NCBI Sequence Read Archive (SRA) under BioProject PRJNA1359397: the Syn1.0 Illumina RNA-seq (accessions SRR35996296, SRR35996297 and SRR35996298), the Syn1.0 PacBio Iso-Seq (SRR36012641, SRR36012642 and SRR36012643), and the Syn3A Oxford Nanopore direct-RNA reads (SRR36199724). The Syn3A Illumina RNA-seq reanalysed here was generated by Sandberg *et al.* [10]. The reference genomes are available from NCBI under accessions CP002027.1 (JCVI-syn1.0) and CP016816.2 (JCVI-syn3A). The mass-spectrometry proteomics data are available from the MassIVE repository under accession MSV000099558 (password: 4zzHbHHqqW7Gl2F5). The Cryo-electron tomography data of Syn3A were obtained from the Cryo-ET Data Portal [42], dataset 10442 [43] (https://cryoetdataportal.czscience.com/datasets/10442). Analysis outputs and the visualization Jupyter Notebook are deposited at Zenodo with link https://zenodo.org/records/22553688. Processed per-operon and per-gene tables and the library quality-control reports are provided as Supplementary Data S1–S4.

## Code availability

All analysis code for this study, covering transcriptome read processing, operon segmentation and annotation, RNA-processing and ribonuclease analysis, transcriptome–proteome correlation, the genome-reduction comparison, and visualization Jupyter Notebook is available at https://github.com/Luthey-Schulten-Lab/ JCVI-Syn1_Syn3A_Transcriptomics_Proteomics.

## Author contributions

Z.L.-S., and J.I.G. conceived and designed the project. A.P.M. advised J.E.C. and Y.-l.G. for the sample preparation of Syn1.0 Illumina and PacBio RNA-sequencing data. S.A.G., K.G., G.J., T.M. and S.S. generated the Syn1.0 and Syn3A Oxford Nanopore direct-RNA sequencing data. Y.Y. performed the proteomics experiments and analysed the proteomic data. E.F., T.A.B., C.J.F. and A.T.A. developed the methodology. E.F. and T.A.B analyzed the data. E.F., T.A.B., J.E.C., Y.Y., S.A.G., C.J.F., J.I.G. and Z.L.-S wrote the original draft. E.F., T.A.B., Z.R.T., A.T.A., J.I.G, and Z.L.-S edited the paper. All authors reviewed the paper.

## Competing interests

APM: The content is solely the responsibility of the authors and does not necessarily represent the official views of the National Institutes of Health.

The remaining authors declare that the research was conducted in the absence of any commercial or financial relationships that could be construed as a potential conflict of interest.

## Supplementary Information

### Supplementary Data

The supplementary items below hold the per-gene and per-operon tables underlying the main figures, together with the raw RNA-sample quality-control reports.

**Supplementary Data S1 (**operon.xlsx). The 459 Syn1.0 operons annotated from PacBio long reads, one row per operon: chromosomal boundaries, member genes and their read coverage, transcription start and termination sites, the *−*10 and *−*35 promoter and intrinsic-terminator signatures, and macromolecular-complex annotation.

**Supplementary Data S2 (**syn1 omics.xlsx). The paired transcriptome and proteome table for all 911 Syn1.0 genes: locus tag, gene name, RNA type, gene product, predicted protein localization, Illumina and PacBio TPM, relative protein abundance (iPM) and absolute copy number, predicted translation initiation rate, and codon-adaptation index.

**Supplementary Data S3 (**genome reduction.xlsx). The genome-reduction analysis, one sheet per level: the 95 deletions with their junction taxonomy (Deletions), the per-operon truncation and gene-deletion classification (Operon classification), the Syn3A-read co-transcription evidence for every tested gene pair (Coexpression), a master per-gene table of relative transcript and protein abundance, fold changes, protein-to-transcript ratio, transcriptional-impact class and curated function with deleted loci flagged (Gene table), the per-function pool-share changes (Function categories), and the deleted-gene occupancy and macromolecular-complex abundances (Summary stats).

**Supplementary Data S4 (quality control).** The raw RNA-sample quality-control reports for the in-house sequencing libraries: Syn1.0 PacBio (cDNA and library preparation), Syn1.0 Illumina (two preparations), Syn1.0 Oxford Nanopore direct-RNA, and Syn3A Oxford Nanopore direct-RNA. The Syn1.0 Illumina samples were quantified with the Qubit RNA High Sensitivity assay, depleted of ribosomal RNA with the Invitrogen RiboMinus^TM^ Bacteria 2.0 Transcriptome Isolation Kit, and monitored for loss of the rRNA peaks on an Agilent 5200 Fragment Analyzer; the Syn1.0 PacBio RNA sample was quantified and rRNA-depleted in the same way, and its cDNA and final libraries were additionally sized on an Agilent Femto Pulse. The direct-RNA samples were rRNA-depleted to below a *≤*3% threshold (loss of the 16S and 23S peaks): the Syn1.0 ONT sample with two rounds of RiboMinus^TM^ Bacteria 2.0 (traced on an Agilent Bioanalyzer 2100; 9.8% then 0.8% rRNA), and the Syn3A ONT sample with a NEBNext® rRNA Depletion Kit for Bacteria (traced on an Agilent TapeStation 2200). Each report is a separate file, provided together as a single archive indexed by a README.

## References

[1] Lachance, JC, Rodrigue, S, Palsson, BO (2019) Minimal cells, maximal knowledge. eLife 8:e45379.

[2] Breuer, M, Earnest, TM, Merryman, C, Wise, KS, Sun, L, Lynott, MR, Hutchison, CA, Smith, HO, Lapek, JD, Gonzalez, DJ, De Crécy-Lagard, V, Haas, D, Hanson, AD, Labhsetwar, P, Glass, JI, Luthey-Schulten, Z (2019) Essential metabolism for a minimal cell. eLife 8:e36842.

[3] Gibson, DG, Glass, JI, Lartigue, C, Noskov, VN, Chuang, RY, Algire, MA, Benders, GA, Montague, MG, Ma, L, Moodie, MM, Merryman, C, Vashee, S, Krishnakumar, R, Assad-Garcia, N, Andrews-Pfannkoch, C, Denisova, EA, Young, L, Qi, ZQ, Segall-Shapiro, TH, Calvey, CH, Parmar, PP, Hutchison, CA, Smith, HO, Venter, JC (2010) Creation of a Bacterial Cell Controlled by a Chemically Synthesized Genome. Science 329:52–56.

[4] Hutchison, CA, Chuang, RY, Vladimir N. Noskov, Assad-Garcia, N, Deerinck, TJ, Ellisman, MH, Gill, J, Kannan, K, Karas, BJ, Ma, L, Pelletier, JF, Qi, ZQ, Richter, RA, Strychalski, EA, Sun, L, Suzuki, Y, Tsvetanova, B, Wise, KS, Smith, HO, Glass, JI, Merryman, C, Gibson, DG, Venter, JC (2016) Design and synthesis of a minimal bacterial genome. Science 351:aad6253.

[5] Pelletier, JF, Sun, L, Wise, KS, Assad-Garcia, N, Karas, BJ, Deerinck, TJ, Ellisman, MH, Mershin, A, Gershenfeld, N, Chuang, RY, Glass, JI, Strychalski, EA (2021) Genetic requirements for cell division in a genomically minimal cell. Cell 184:2430–2440.e16.

[6] Thornburg, ZR, Maytin, A, Kwon, J, Brier, TA, Gilbert, BR, Fu, E, Gao, YL, Quenneville, J, Wu, T, Li, H, Long, T, Pezeshkian, W, Sun, L, Bittencourt, DMdC, Glass, JI, Mehta, AP, Ha, T, Luthey-Schulten, Z (2026) Bringing the genetically minimal cell to life on a computer in 4D. Cell 189:2582–2597.e27.

[7] Bittencourt, DMDC, Brown, DM, Assad-Garcia, N, Romero, MR, Sun, L, Palhares De Melo, LAM, Freire, M, Glass, JI (2024) Minimal Bacterial Cell JCVI-syn3B as a Chassis to Investigate Interactions between Bacteria and Mammalian Cells. ACS Synth. Biol. 13:1128–1141.

[8] Gilbert, BR, Thornburg, ZR, Lam, V, Rashid, FZM, Glass, JI, Villa, E, Dame, RT, Luthey-Schulten, Z (2021) Generating Chromosome Geometries in a Minimal Cell From Cryo-Electron Tomograms and Chromosome Conformation Capture Maps. Front. Mol. Biosci. 8:644133.

[9] Justice, I, Kiesel, P, Safronova, N, von Appen, A, Saenz, JP (2024) A tuneable minimal cell membrane reveals that two lipid species suffice for life. Nat Commun 15:9679.

[10] Sandberg, TE, Wise, KS, Dalldorf, C, Szubin, R, Feist, AM, Glass, JI, Palsson, BO (2023) Adaptive evolution of a minimal organism with a synthetic genome. iScience 26:107500.

[11] Moger-Reischer, RZ, Glass, JI, Wise, KS, Sun, L, Bittencourt, DMC, Lehmkuhl, BK, Schoolmaster, DR, Lynch, M, Lennon, JT (2023) Evolution of a minimal cell. Nature 620:122–127.

[12] Thornburg, ZR, Bianchi, DM, Brier, TA, Gilbert, BR, Earnest, TM, Melo, MC, Safronova, N, Saénz, JP, Cook, AT, Wise, KS, Hutchison, CA, Smith, HO, Glass, JI, Luthey-Schulten, Z (2022) Fundamental behaviors emerge from simulations of a living minimal cell. Cell 185:345–360.e28.

[13] Razin, S, Yogev, D, Naot, Y (1998) Molecular Biology and Pathogenicity of Mycoplasmas. Microbiol Mol Biol Rev 62:1094–1156.

[14] Jacob, F, Monod, J (1961) Genetic regulatory mechanisms in the synthesis of proteins. Journal of Molecular Biology 3:318–356.

[15] Stark, R, Grzelak, M, Hadfield, J (2019) RNA sequencing: the teenage years. Nat Rev Genet 20:631–656.

[16] Yan, B, Boitano, M, Clark, TA, Ettwiller, L (2018) SMRT-Cappable-seq reveals complex operon variants in bacteria. Nat Commun 9:3676.

[17] Grünberger, F, Ferreira-Cerca, S, Grohmann, D (2022) Nanopore sequencing of RNA and cDNA molecules in Escherichia coli. RNA 28:400–417.

[18] Ju, X, Li, D, Liu, S (2019) Full-length RNA profiling reveals pervasive bidirectional transcription terminators in bacteria. Nat Microbiol 4:1907–1918.

[19] Mondal, S, Yakhnin, AV, Sebastian, A, Albert, I, Babitzke, P (2016) NusA-dependent transcription termination prevents misregulation of global gene expression. Nat Microbiol 1:15007.

[20] Mandell, ZF, Vishwakarma, RK, Yakhnin, H, Murakami, KS, Kashlev, M, Babitzke, P (2022) Comprehensive transcription terminator atlas for Bacillus subtilis. Nat Microbiol 7:1918–1931.

[21] Lloréns-Rico, V, Lluch-Senar, M, Serrano, L (2015) Distinguishing between productive and abortive promoters using a random forest classifier in Mycoplasma pneumoniae. Nucleic Acids Res 43:3442–3453.

[22] Matteau, D, Lachance, J, Grenier, F, Gauthier, S, Daubenspeck, JM, Dybvig, K, Garneau, D, Knight, TF, Jacques, P, Rodrigue, S (2020) Integrative characterization of the near-minimal bacterium Mesoplasma florum. Molecular Systems Biology 16:e9844.

[23] Kingsford, CL, Ayanbule, K, Salzberg, SL (2007) Rapid, accurate, computational discovery of Rho-independent transcription terminators illuminates their relationship to DNA uptake. Genome Biol 8:R22.

[24] Pelletier, JF, Glass, JI, Strychalski, EA (2022) Cellular mechanics during division of a genomically minimal cell. Trends in Cell Biology 32:900–907.

[25] Kaźmierczak, J, Kremer, B (2019) Pattern of cell division in 3.4Ga-old microbes from South Africa. Precambrian Research 331:105357.

[26] Durand, S, Condon, C (2018) RNases and Helicases in Gram-Positive Bacteria. Microbiol Spectr 6:6.2.16.

[27] Huch, S, Nersisyan, L, Ropat, M, Barrett, D, Wu, M, Wang, J, Valeriano, VD, Vardazaryan, N, Huerta-Cepas, J, Wei, W, Du, J, Steinmetz, LM, Engstrand, L, Pelechano, V (2023) Atlas of mRNA translation and decay for bacteria. Nat Microbiol 8:1123–1136.

[28] Janssen, BD, Hayes, CS (2012) The tmRNA ribosome rescue system. Adv Protein Chem Struct Biol 86:151–191.

[29] Fu, E, Thornburg, ZR, Brier, TA, Wei, R, Yuan, B, Gilbert, BR, Wang, S, Luthey-Schulten, Z (2026) Assembly of Macromolecular Complexes in the Whole-Cell Model of a Minimal Cell. J. Phys. Chem. B 130:11–32.

[30] Taggart, J, Dierksheide, K, LeBlanc, H, Lalanne, JB, Durand, S, Braun, F, Condon, C, Li, GW (2025) A high-resolution view of RNA endonuclease cleavage in Bacillus subtilis. Nucleic Acids Res 53:gkaf030.

[31] Roots, CT, Lukasiewicz, A, Barrick, JE (2021) OSTIR: open source translation initiation rate prediction. J Open Source Softw 6:3362.

[32] Li, GW, Burkhardt, D, Gross, C, Weissman, J (2014) Quantifying Absolute Protein Synthesis Rates Reveals Principles Underlying Allocation of Cellular Resources. Cell 157:624–635.

[33] Sharp, PM, Li, WH (1987) The codon adaptation index-a measure of directional synonymous codon usage bias, and its potential applications. Nucleic Acids Res 15:1281–1295.

[34] Razin, S, Argaman, M, Avigan, J (1963) Chemical Composition of Mycoplasma Cells and Membranes. Microbiology 33:477–487.

[35] Bratbak, G, Dundas, I (1984) Bacterial dry matter content and biomass estimations. Appl Environ Microbiol 48:755–757.

[36] Leblanc, G, Le Grimellec, C (1979) Active K+ transport in *Mycoplasma mycoides* var. Capri. Net and unidirectional K+ movements. Biochimica et Biophysica Acta (BBA) - Biomembranes 554:156–167.

[37] Lloréns-Rico, V, Cano, J, Kamminga, T, Gil, R, Latorre, A, Chen, WH, Bork, P, Glass, JI, Serrano, L, Lluch-Senar, M (2016) Bacterial antisense RNAs are mainly the product of transcriptional noise. Science Advances 2:e1501363.

[38] Redko, Y, Aubert, S, Stachowicz, A, Lenormand, P, Namane, A, Darfeuille, F, Thibonnier, M, De Reuse, H (2013) A minimal bacterial RNase J-based degradosome is associated with translating ribosomes. Nucleic Acids Research 41:288–301.

[39] Earnest, T, Lai, J, Chen, K, Hallock, M, Williamson, J, Luthey-Schulten, Z (2015) Toward a Whole-Cell Model of Ribosome Biogenesis: Kinetic Modeling of SSU Assembly. Biophysical Journal 109:1117–1135.

[40] Earnest, TM, Cole, JA, Peterson, JR, Hallock, MJ, Kuhlman, TE, Luthey-Schulten, Z (2016) Ribosome biogenesis in replicating cells: Integration of experiment and theory. Biopolymers 105:735–751 eprint: https://onlinelibrary.wiley.com/doi/pdf/10.1002/bip.22892.

[41] Davis, JH, Tan, YZ, Carragher, B, Potter, CS, Lyumkis, D, Williamson, JR (2016) Modular Assembly of the Bacterial Large Ribosomal Subunit. Cell 167:1610–1622.e15.

[42] Ermel, U, Cheng, A, Ni, JX, Gadling, J, Venkatakrishnan, M, Evans, K, Asuncion, J, Sweet, A, Pourroy, J, Wang, ZS, Khandwala, K, Nelson, B, McCarthy, D, Wang, EM, Agarwal, R, Carragher, B (2024) A data portal for providing standardized annotations for cryo-electron tomography. Nat Methods 21:2200–2202.

[43] All, M, Arlana, P, Yue, Y, Jonathan, S, Elizabeth, M, John, G, Shawn, Mohammadreza, P (2024) CryoET of near-minimal cells Mycoplasma mycoides JCVI-Syn3A for the development of subtomogram averaging pipelines.

[44] Maier, T, Schmidt, A, Güell, M, Kühner, S, Gavin, A, Aebersold, R, Serrano, L (2011) Quantification of mRNA and protein and integration with protein turnover in a bacterium. Mol Syst Biol 7:MSB201138.

[45] Ishihama, Y, Schmidt, T, Rappsilber, J, Mann, M, Hartl, FU, Kerner, MJ, Frishman, D (2008) Protein abundance profiling of the Escherichia coli cytosol. BMC Genomics 9:102.

[46] Ingolia, NT, Ghaemmaghami, S, Newman, JRS, Weissman, JS (2009) Genome-Wide Analysis in Vivo of Translation with Nucleotide Resolution Using Ribosome Profiling. Science 324:218–223.

[47] Michalik, S, Reder, A, Richts, B, Faßhauer, P, Mäder, U, Pedreira, T, Poehlein, A, van Heel, AJ, van Tilburg, AY, Altenbuchner, J, Klewing, A, Reuß, DR, Daniel, R, Commichau, FM, Kuipers, OP, Hamoen, LW, Völker, U, Stülke, J (2021) The Bacillus subtilis Minimal Genome Compendium. ACS Synth. Biol. 10:2767–2771.

[48] Taggart, JC, Lalanne, JB, Li, GW (2021) Quantitative Control for Stoichiometric Protein Synthesis. Annu. Rev. Microbiol. 75:243–267.

[49] Mortazavi, A, Williams, BA, McCue, K, Schaeffer, L, Wold, B (2008) Mapping and quantifying mammalian transcriptomes by RNA-Seq. Nat Methods 5:621–628.

[50] Wenger, AM, Peluso, P, Rowell, WJ, Chang, PC, Hall, RJ, Concepcion, GT, Ebler, J, Fungtammasan, A, Kolesnikov, A, Olson, ND, Töpfer, A, Alonge, M, Mahmoud, M, Qian, Y, Chin, CS, Phillippy, AM, Schatz, MC, Myers, G, DePristo, MA, Ruan, J, Marschall, T, Sedlazeck, FJ, Zook, JM, Li, H, Koren, S, Carroll, A, Rank, DR, Hunkapiller, MW (2019) Accurate circular consensus long-read sequencing improves variant detection and assembly of a human genome. Nat Biotechnol 37:1155–1162.

[51] Rhoads, A, Au, KF (2015) PacBio Sequencing and its Applications. genom. proteom. bioinform. 13:278–289.

[52] Garalde, DR, Snell, EA, Jachimowicz, D, Sipos, B, Lloyd, JH, Bruce, M, Pantic, N, Admassu, T, James, P, Warland, A, Jordan, M, Ciccone, J, Serra, S, Keenan, J, Martin, S, McNeill, L, Wallace, EJ, Jayasinghe, L, Wright, C, Blasco, J, Young, S, Brocklebank, D, Juul, S, Clarke, J, Heron, AJ, Turner, DJ (2018) Highly parallel direct RNA sequencing on an array of nanopores. Nat Methods 15:201–206.

[53] Soneson, C, Yao, Y, Bratus-Neuenschwander, A, Patrignani, A, Robinson, MD, Hussain, S (2019) A comprehensive examination of Nanopore native RNA sequencing for characterization of complex transcriptomes. Nat Commun 10:3359.

[54] Liu, H, Begik, O, Lucas, MC, Ramirez, JM, Mason, CE, Wiener, D, Schwartz, S, Mattick, JS, Smith, MA, Novoa, EM (2019) Accurate detection of m6A RNA modifications in native RNA sequences. Nat Commun 10:4079.

[55] Grünberger, F, Reichelt, R, Bunk, B, Spröer, C, Overmann, J, Rachel, R, Grohmann, D, Hausner, W (2019) Next Generation DNA-Seq and Differential RNA-Seq Allow Re-annotation of the Pyrococcus furiosus DSM 3638 Genome and Provide Insights Into Archaeal Antisense Transcription. Front. Microbiol. 10.

[56] Langmead, B, Salzberg, SL (2012) Fast gapped-read alignment with Bowtie 2. Nat Methods 9:357–359.

[57] Li, H, Handsaker, B, Wysoker, A, Fennell, T, Ruan, J, Homer, N, Marth, G, Abecasis, G, Durbin, R, 1000 Genome Project Data Processing Subgroup (2009) The Sequence Alignment/Map format and SAMtools. Bioinformatics 25:2078–2079.

[58] Li, H (2018) Minimap2: pairwise alignment for nucleotide sequences. Bioinformatics 34:3094–3100.

[59] Martin, KR, Le, HT, Abdelgawad, A, Yang, C, Lu, G, Keffer, JL, Zhang, X, Zhuang, Z, Asare-Okai, PN, Chan, CS, Batish, M, Yu, Y (2024) Development of an efficient, effective, and economical technology for proteome analysis. Cell Reports Methods 4:100796.

[60] Sun, J, Xu, X, Wei, S, Yu, Y (2026) In-Cell Proteomics Enables High-Resolution Spatial and Temporal Mapping of Early *Xenopus tropicalis* Embryos. Molecular & Cellular Proteomics 25:101481.

[61] Milo, R (2013) What is the total number of protein molecules per cell volume? A call to rethink some published values. BioEssays 35:1050–1055 eprint: https://onlinelibrary.wiley.com/doi/pdf/10.1002/bies.201300066.

[62] Hallgren, J, Tsirigos, KD, Pedersen, MD, Almagro Armenteros, JJ, Marcatili, P, Nielsen, H, Krogh, A, Winther, O (2022) DeepTMHMM predicts alpha and beta transmembrane proteins using deep neural networks.

[63] Teufel, F, Almagro Armenteros, JJ, Johansen, AR, Gíslason, MH, Pihl, SI, Tsirigos, KD, Winther, O, Brunak, S, Von Heijne, G, Nielsen, H (2022) SignalP 6.0 predicts all five types of signal peptides using protein language models. Nat Biotechnol 40:1023–1025.

[64] Kurtz, S, Phillippy, A, Delcher, AL, Smoot, M, Shumway, M, Antonescu, C, Salzberg, SL (2004) Versatile and open software for comparing large genomes. Genome Biol 5:R12.

[65] Quinlan, AR, Hall, IM (2010) BEDTools: a flexible suite of utilities for comparing genomic features. Bioinformatics 26:841–842.

